# High-resolution temporal profiling reveals synchronized dynamics of the mouse gut microbiome

**DOI:** 10.64898/2026.03.26.714232

**Authors:** Rina Kurokawa, Rie Maskawa, Musashi Arakawa, Hiroaki Masuoka, Hideki Takayasu, Yu Yoshikawa, Tanzila Raihan, Chie Shindo, Kaoru Kaida, Misa Takagi, Maki Tanokura, Iwijn De Vlaminck, Lena Takayasu, Misako Takayasu, Wataru Suda

**Author notes:** Corresponding authors: Wataru Suda., Misako Takayasu., Lena Takayasu. These authors contributed equally to this work (Rina Kurokawa, Rie Maskawa).

## Abstract

**Background:** The gut microbiome undergoes diurnal oscillations entrained by host circadian rhythms and feeding behavior, and disruption of these oscillations has been linked to metabolic dysfunction. However, the temporal organization of microbial activity within these oscillatory cycles remains poorly understood, largely because conventional sampling approaches lack the temporal resolution needed to resolve rapid microbial dynamics.

**Results:** To address this gap, we developed a fully automated fecal collection system that enables continuous sampling from individual mice at minute- to hour-scale resolution without manual intervention. Using this platform, we collected fecal samples at hourly intervals over 14 consecutive days from individually housed mice. To reconstruct genome- and function-level trajectories from these data, we developed a cost-effective method (GORIN) that combines full-length 16S rRNA gene amplicon sequencing with long-read metagenomic references. Applying a newly developed phase-analytic method (NPPD) to determine the instantaneous phase of each genome and functional gene, we found that the gut microbiota is organized into two phase-coherent groups corresponding to the light and dark phases of the host circadian cycle. These two groups differed in gene content, suggesting distinct carbohydrate utilization strategies, with the light-phase community enriched in mucosal glycan-degrading enzymes, whereas the dark-phase community was enriched in carbon storage metabolism and motility-related functions. Within each phase, microbial taxa and functional genes displayed reproducible temporal succession across individual mice. Perturbations, including cage transfer and antibiotic treatment transiently disrupted phase synchronization, as measured by the phase order parameter, followed by gradual recovery of phase coherence.

**Conclusions:** These findings reveal reproducible phase-synchronized temporal organization in the mouse gut microbiome, with community-level phase relationships that can be quantified and tracked over time and following perturbation. The automated sampling platform, genome-resolved profiling framework, and phase-analytic metrics introduced here provide a scalable strategy for characterizing community-level temporal dynamics in host-associated microbial ecosystems, with potential applications for understanding how disruption of microbial temporal organization contributes to dysbiosis

## Background

The gut microbiome is a highly dynamic ecosystem whose composition and activity fluctuate on timescales ranging from hours to days [1–5]. A growing body of evidence has established that these temporal fluctuations are not random noise but are organized by host circadian rhythms and feeding behavior. Pioneering studies demonstrated that gut microbial communities undergo diurnal oscillations entrained by host feeding patterns, and that disruption of host clock genes or feeding rhythmicity abolishes these oscillations, leading to dysbiosis and metabolic dysfunction [1–4]. Subsequent work revealed that microbial metabolites, particularly short-chain fatty acids, feedback to modulate host circadian clock gene expression in intestinal epithelial cells and hepatocytes [6], and that the microbiota programs diurnal rhythms in host metabolism through epigenetic mechanisms such as histone deacetylase 3 [7]. These findings establish a bidirectional circadian dialogue between the host and its microbiota and demonstrate the functional relevance of microbiome rhythmicity to host physiology.

Despite these advances in understanding the functional significance of microbiome rhythmicity, the temporal organization of microbial taxa and functional gene content within these oscillatory cycles, including their sequential succession within the community, remains poorly understood. Resolving such fine-scale organization requires continuous observation at high temporal resolution over extended periods, together with genome- and function-resolved profiling. Most studies, however, have relied on sparse sampling of 4 to 8 time points per 24-hour cycle [1,4,6,8], reflecting the lack of platforms for continuous, high-resolution collection: conventional fecal sampling requires manual intervention at each time point, and the handling-induced stress it entails can itself alter the microbiota, confounding the very dynamics under study [9]. Genome- and function-resolved profiling of densely sampled series is further constrained by cost and throughput, since shotgun metagenomics is cost-prohibitive at hundreds or thousands of time points, while amplicon sequencing, though inexpensive and high-throughput, lacks the required functional resolution.

To overcome these limitations, we developed a fully automated fecal collection system for mice that enables continuous sampling at minute-to hour-scale resolution over weeks without manual intervention or animal handling, thereby preserving the host’s natural behavioral and physiological context. To recover genome- and function-resolved temporal trajectories from the resulting densely sampled series, we developed GORIN (Genome abundance Optimization from 16S rRNA-Informed Networks), which integrates full-length 16S rRNA gene amplicon profiles with a limited set of long-read metagenomic references. This approach enables cost-effective profiling of genome- and function-level dynamics across hundreds of time points.

To characterize this unresolved temporal organization, we focus on the relative timing of oscillations across taxa, which can be represented in terms of phase [10]. A phase representation provides a common coordinate for comparing relative timing across oscillatory components despite differences in amplitude and waveform, and has been widely used to characterize temporal relationships in other biological oscillatory systems, including neural signals [11] and oscillating cell populations [12]. We therefore developed NPPD (Non-parametric Period Detection), a phase-analytic framework for determining the instantaneous phase of individual microbial and functional trajectories.

In this study, we combined automated hourly sampling over 14 consecutive days with GORIN and NPPD to resolve the fine-scale temporal organization of microbial genomes and functions in the mouse gut microbiota. We described the resulting phase structure across taxa and functional genes, quantified its coherence over time, and examined how phase organization changed following environmental and antibiotic perturbations. Our findings provide a high-resolution description of community-level phase organization in the gut microbiome and define the temporal structure that future mechanistic studies will need to explain.

## Methods

### Animal experiments and fecal sampling

All animal experiments were approved by the Animal Experiments Committee of RIKEN (approval number AEY2024-32). Female C57BL/6J mice were purchased from CLEA Japan, Inc. and acclimated for more than 1 week in the RIKEN animal facility under specific-pathogen-free (SPF) conditions. During acclimation, mice were group-housed under temperature- and humidity-controlled conditions with a 12-h light/dark cycle (lights on at 07:00, defined as Zeitgeber time 0 (ZT0)), and provided *ad libitum* access to a standard diet (CLEA Rodent Diet CE-2; CLEA Japan) and water.

At the time of experimentation, mice were 10–22 weeks old. Detailed information on the individual mice and experimental schedules is provided in Additional file 2: Table S1. Six mice (A–F) were used for the time-series experiments. Mouse A and Mouse B were sampled without intervention for 14 days. Mouse C and Mouse D were sampled for 18 days as the control and vancomycin-treated animals, respectively, in the antibiotic perturbation experiment described below. Mouse E and Mouse F were sampled for 4 days each. Samples from all six mice were used to reconstruct genome- and function-level temporal dynamics and examine phase organization. Six additional mice (V1–V4, P1, and P2) were used to validate the collection system; samples from Mouse E and Mouse F were also used for this validation. During the experiments, mice were housed individually in metabolic cages that constituted part of an automated mouse fecal collection system, maintained under the same temperature, humidity, and light–dark cycle conditions, with *ad libitum* access to the same diet and water.

Fecal pellets were automatically collected into tubes pre-filled with RNA*later™* solution (Thermo Fisher Scientific Inc., Waltham, MA, USA) at 1-h intervals unless otherwise specified and stored at 4 °C until further processing.

For antibiotic perturbation experiments, vancomycin hydrochloride (biochemistry grade; FUJIFILM Wako Pure Chemical Corp., Osaka, Japan) was dissolved in drinking water at a concentration of 2.5 mg/mL and sterilized using a 0.22 µm filter. The solution (200 µL per mouse) was administered once via oral gavage at 15:00 on day 4 after the initiation of automated fecal collection. Mouse D received the vancomycin solution, whereas Mouse C (control) received an equivalent volume of sterile drinking water under identical conditions.

To assess sampling at shorter intervals, Mouse P1 and Mouse P2 were sampled at 1-h intervals for the first 3 days, followed on day 4 by collection of 24 samples at 10-min and 5-min intervals, respectively.

To validate the automated collection workflow against manual collection followed by immediate DNA extraction, paired sampling was performed in four additional mice (V1–V4) at 14:00 (ZT7) and 19:00 (ZT12). Immediately after confirming that a fecal pellet had been excreted and collected by the automated system, the mouse was taken out of the device, and a naturally excreted pellet was collected manually within one hour of automated collection. Manually collected pellets were placed in RNA*later*™ and processed immediately for DNA extraction. Automatically collected pellets were retained in RNA*later*™ in the device and retrieved for DNA extraction 24 h after collection, following the routine operating procedure. This yielded eight matched pairs.

To contextualize differences between the collection workflows, hourly samples from Mouse E, Mouse F, Mouse P1, and Mouse P2 were used to assess within- and between-mouse variation. Samples were selected from the 24-h period immediately following the first 48 h of collection, yielding 21, 19, 21, and 21 samples, respectively. Details of the validation analyses are provided in Additional file 1: Supplementary Notes.

### DNA extraction

Bacterial genomic DNA was extracted from mouse fecal samples preserved in RNA*later*™ solution (Thermo Fisher Scientific Inc., Waltham, MA, USA) using an enzymatic lysis method with modifications [13]. A single fecal pellet was retrieved from the storage tube, briefly blotted to remove residual RNA*later*™, and weighed. The sample was manually homogenized in phosphate-buffered saline (PBS; pH 7.4) using the tip of a pipette and washed once with PBS by centrifugation at 6,000 × *g* for 10 min at 4 °C. The pellet was resuspended in 10 mM Tris–20 mM EDTA (TE20).

Microbial cells were lysed with 15 mg lysozyme (Sigma-Aldrich Co., St. Louis, MO, USA) and 2,000 U achromopeptidase (Fujifilm Wako Pure Chemical Corp., Osaka, Japan) at 37 °C for 2 h, followed by incubation with 1% sodium dodecyl sulfate (Fujifilm Wako Pure Chemical Corp.) and 1 mg proteinase K (Merck KGaA, Darmstadt, Germany) at 55 °C for 1 h. Lysates were mixed with an equal volume of phenol–chloroform–isoamyl alcohol (25:24:1; Nippon Gene Co. Ltd., Tokyo, Japan) for 10 min at 24 °C, and the aqueous phase was recovered by centrifugation.

Sodium acetate (3 M; 10% v/v) was added to the supernatant, and DNA was precipitated with isopropanol. The DNA pellet was collected by centrifugation at 12,000 × *g* for 10 min at 4 °C, rinsed once with 75% ethanol, and dissolved in 100 µL of 10 mM Tris–1 mM EDTA (TE). The solution was treated with RNase A (10 mg mL^−1^; Nippon Gene Co. Ltd.) for 30 min at 37 °C, then mixed with 0.6 volumes of 20% polyethylene glycol 6,000 and 2.5 M sodium chloride (Hampton Research Corp., Aliso Viejo, USA) and incubated on ice for ≥ 10 min. DNA was pelleted again at 12,000 × *g* for 10 min at 4 °C, washed twice with 75% ethanol, and dissolved in 50 µL TE.

### Full-length 16S rRNA gene amplicon sequencing

Full-length 16S rRNA genes were amplified using the primers PacBio Kinnex barcoded forward primer: (5′-CTAC ACGA CGCT CTTC CGAT CT NNNNNNNNNN AGRG TTYG ATYM TGGC TCAG-3′) and PacBio Kinnex barcoded 1492Rmod: (5′-AAGC AGTG GTAT CAAC GCAG AG NNNNNNNNNN RGHT ACCT TGTT ACGA CTT-3′), where “N” denotes a unique PacBio Kinnex barcode sequence. Subsequent procedures followed the manufacturer’s protocol, “Preparing Kinnex™ libraries from 16S rRNA amplicons” (PacBio, Pacific Biosciences of California, Inc.).

The sequencing library was prepared using the PacBio SMRTbell® Prep Kit 3.0 according to the manufacturer’s protocol. Sequencing was performed on the PacBio Revio™ system with the option Full resolution base qual = TRUE. PacBio HiFi reads were generated automatically using SMRT Link software [14] (v25.1) with default parameters. Read segmentation was conducted using Skera (v1.3.0), and demultiplexing was performed with Lima (v2.12.0) under the HIFI-ASYMMETRIC preset.

### Full-length 16S rRNA gene amplicon sequence variants analysis

Full-length 16S rRNA gene amplicon sequence variants (ASVs) were inferred from demultiplexed HiFi reads using the DADA2 package [15] (v1.30.0) in R (v4.3.3) according to the previously described DADA2 for PacBio workflow [16] with slight modifications. The reads were subjected to quality filtering and trimming using the filterAndTrim function with the following parameters: minQ = 3, minLen = 1300, maxLen = 1600, maxN = 0, rm.phix = FALSE, maxEE = 5. ASVs were then subjected to a homology search against SBDI Sativa curated 16S GTDB database [17] (v10) using BLASTN [18] (v2.16+), with a maximum E-value cut-off of 1 × 10^-10^. Top hits were determined by the highest bitscore. We quantified community composition using a spike-in standard by adding a fixed amount of genomic DNA from *Salinibacter ruber* to each fecal DNA sample. The observed copy number of the *S. ruber* standard was then used to convert ASV abundances to absolute values, reported as copies per milligram of feces.

### Long-read shotgun metagenomic sequencing

For long-read shotgun metagenomic sequencing, four composite DNA pools were prepared for each of Mouse A–D, separating samples by light phase (ZT0–11), dark phase (ZT12–23), and collection period. For Mouse E and Mouse F, samples from all time points over the four-day collection period were combined into a single pool per mouse. Equal amounts of DNA from the time points assigned to each pool were combined. The number of samples, first and last sampling days, and light/dark phase represented in each pool are provided in Additional file 2: Table S2. The sequencing library for each pooled sample was prepared using the PacBio SMRTbell® Prep Kit 3.0 following the manufacturer’s protocol. Sequencing was performed on the PacBio Revio™ system with the option Full resolution base qual = TRUE. HiFi reads were generated automatically using SMRT Link software (v25.1) with default parameters.

### Construction of long-read metagenome-assembled genomes

For long-read shotgun metagenomic data, reads mapped to internal control sequences or host genomes were removed using minimap2 [19] (v2.28-r1209), with an identity threshold of ≥ 90 % and alignment coverage of ≥ 85 %.

De novo assemblies were generated independently with hifiasm-meta [20] (v0.25), metaFlye [21] (v2.9), HiCanu [22] (v2.3), metaMDBG [23] (v1.1), and myloasm [24] (v0.3.0). Contigs obtained from each assembler were binned separately with SemiBin2 [25] (v2.2.0) to recover metagenome-assembled genomes (MAGs).

MAG quality was assessed using CheckM2 [26] (v1.1.0) for genome completeness and contamination, and potential chimeric bins were further screened with GUNC [27] (v1.0.5). MAGs with ≥ 50 % completeness and ≤ 10 % contamination were designated as high- and medium-quality metagenome-assembled genomes (HMQ-MAGs). Taxonomic classification was performed using GTDB-Tk [28] (v2.3.2, GTDB release R214).

### Construction of correspondence matrices

ASV sequences were searched against HMQ-MAGs using BLASTN (v2.16.0+), and only hits with 100 % nucleotide identity and 100 % alignment coverage were retained. Based on these exact matches, an ASV–MAG correspondence matrix was constructed, with MAGs represented as columns and ASVs as rows, summarizing the number and types of ASVs corresponding to each MAG (see Additional file 2: Tables S4–9). For MAGs corresponding to identical ASVs, representative MAGs were selected based on the highest completeness and lowest contamination (see Additional file 2: Table S10; hereafter, referred to as MAGs).

### Functional annotation

Genes were predicted from HMQ-MAGs using Bakta [29–31] (v1.11.3). Functional annotation was performed by assigning Kyoto Encyclopedia of Genes and Genomes (KEGG) [32] orthology using KofamScan [33] (v1.3.0) and identifying carbohydrate-active enzymes (CAZymes) [34] using dbCAN3 [35].

The numbers and types of KEGG orthologs (KOs) and CAZyme families identified in each MAG are summarized in Additional file 2: Tables S11–22.

### Genome abundance Optimization from 16S rRNA-Informed Networks (GORIN)

To infer MAG abundances from ASV counts, we developed Genome abundance Optimization from 16S rRNA-Informed Networks (GORIN), an optimization framework based on an iterative Poisson-weighted nonnegative least squares formulation with a network-based regularization term. GORIN estimates MAG abundances independently at each time point t, accounting for both measurement noise and ASV sharing among MAGs. Let the number of MAG types be *M* and the number of ASV types be *N*. Let *y*_*i*_ ∈ ℤ _≥0_ denote the observed count of ASV *i*, and *C*_*ij*_ represent the copy number of ASV *i* contained in MAG *j*. The abundance of MAG *j* is denoted as *x*_*j*_ ∈ ℝ_≥0_, which is estimated in the following framework.

### Network construction based on ASV–MAG correspondence

To identify potential redundancy in the copy number matrix C and improve numerical stability, we represented the ASV–MAG correspondence as a bipartite network. Specifically, we constructed a network H from the copy number matrix C, where ASVs and MAGs are represented as distinct node types, and edges are assigned between each MAG and the ASVs it contains. Based on the definition from the copy number matrix *C*, when an element *C*_*ij*_ is nonzero, an edge is established between the ASV node *a*_*i*_ and the MAG node *m*_*j*_ corresponding to that element. By extracting connected components from the constructed network *H*, we obtain independent subnetworks that do not share edges. Each subnetwork defines a local system of linear equations that describes the abundance relationship between ASVs and MAGs within that component.

For the *p*-th connected component, let 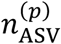 and 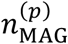 denote the number of ASV and MAG types, respectively. Let 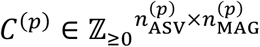 be the corresponding submatrix of *C*.

Depending on the rank *r*^(*p*)^ = rank7(*C*^(*p*)^), the systems are classified as follows:

1. Full-rank linear systems: when 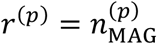,
2. Rank-deficient linear systems: when 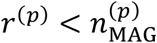.

GORIN handles these two cases differently. For full-rank systems, abundances are estimated by Poisson-based nonnegative iteratively reweighted least squares. For rank-deficient systems, a network-structure-based smoothing term is added to the same formulation to resolve the indeterminacy.

### Nonnegative-constrained Iteratively Reweighted Least Squares Based on the Poisson Process

The expected ASV count *µ*_*i*_ is modeled as a linear combination of the MAG abundances:

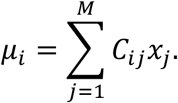

As sequencing read counts are observed as discrete stochastic events, we assume that the observed count of each ASV follows a Poisson process:

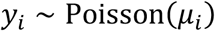

For the Poisson distribution with a linear link function *g*(*µ*) = *µ*, the updates of maximum likelihood estimation can be formulated using IRWLS, with the working variable *y*_*i*_ and the weight 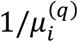 [36]. Accordingly, the estimation reduces to solving the following optimization problem:

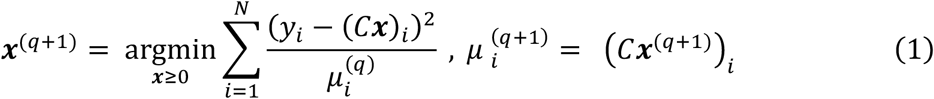

where *q* denotes the iteration number, and the initial value is set as 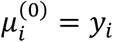.

The nonnegativity constraint ***x*** ≥ 0 is imposed to ensure that the estimated abundance of each MAG remains physically meaningful.

This procedure iteratively approximates the maximization of the Poisson likelihood under the nonnegativity constraint.

### Network-regularized smoothing for underdetermined systems

When 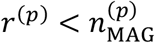, the corresponding system of linear equations becomes underdetermined. To address this indeterminacy, we introduce a network-structure-based smoothing term.

It is difficult to directly determine from which MAG an ASV originated based on the available data. In this study, for such underdetermined systems, we introduce a smoothing penalty that encourages similar abundances among MAGs sharing ASVs, although their true abundances may differ. Furthermore, if MAG *j* and MAG ℓ share *W*_*j*ℓ_ ASVs, we define the connection strength between them as:

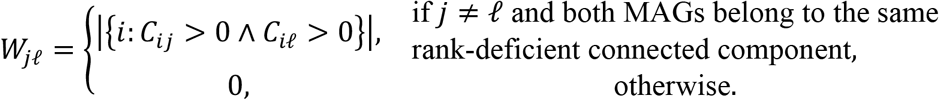

Here, |*S*| denotes the number of elements in the set *S*.

Thus, the smoothing term for each connection is expressed as:

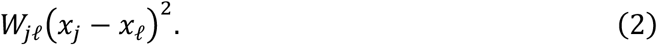

By adding the smoothing term Eq. (2) to Eq. (1), the optimization problem is extended as follows:

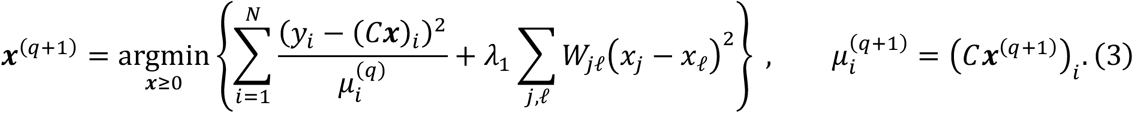

where, *W* = (*W*_*j*ℓ_) denotes the adjacency weight matrix of the network, and the weighted graph Laplacian matrix *L* = (*L*_*j*ℓ_) is defined as:

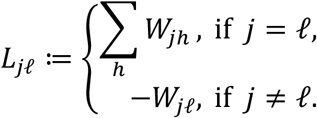

The Laplacian matrix reflects the network structure and quantifies smoothness between connected nodes. The following identity holds [37]:

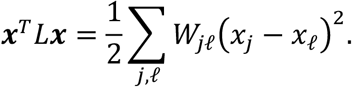

Therefore, defining *λ* = 2*λ*_1_, Eq. (3) can be rewritten as:

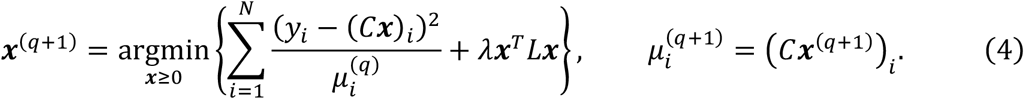

Equation (4) is equivalent to Eq. (1) when all extracted connected components contain no underdetermined subsystems.

### Numerical implementation

In our implementation, two numerical adjustments were applied. First, to avoid division by zero when 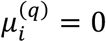 in the term 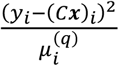, the denominator 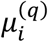 was replaced by max 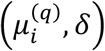, where *δ* > 0 . Second, when network regularization was applied, a small positive value was added to the diagonal elements of *L*, i.e., *L* → *L* + *ϵI* with *ϵ* > 0, to obtain a positive-definite matrix. This diagonal adjustment applies to all MAGs, whereas the smoothing edges in *L* are restricted to rank-deficient components. The optimization problem with these numerical corrections is expressed as:

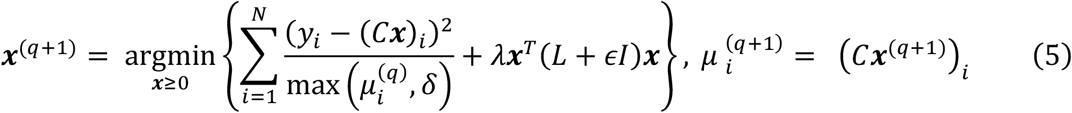

As the matrix *L* + *ϵI* is positive definite, it can be decomposed using Cholesky factorization as

*L* + *ϵI* = *R*^*T*^*R*. Defining 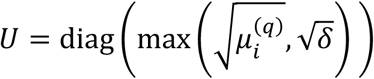, the following extended matrices can be constructed:

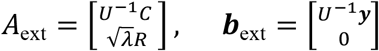 can be rewritten as a standard nonnegative least squares (NNLS) problem:

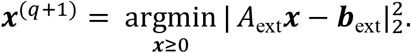

When all connected components are full rank, the regularization term is omitted, and *A*_ext_ = *U*^−1^*C* and ***b***_ext_ = *U*^−1^***y***.

The NNLS problem was solved using the *scipy*.*optimize*.*nnls* function based on the algorithm of Lawson and Hanson [38]. In the iterative computation, the maximum number of iterations was set to 50, and convergence was determined when the relative change of the estimated vector satisfied

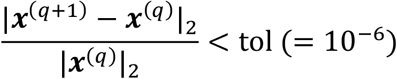

Additionally, in parameter tuning, numerically stable behavior was achieved by setting a sufficiently small *λ* (see Additional file 1: Fig. S1a). In this study, we set *λ* = 1.0 × 10^−8^, *ε* = 1.0 × 10^−4^, and *δ* = 1.0 × 10^−4^. Even when the solution is theoretically unique, numerical estimates may vary with finite-precision arithmetic, stopping criteria, and the conditioning of the problem.

An implementation of the GORIN is publicly available on GitHub at https://github.com/musashi021203/GORIN.

### Assessment of internal consistency in MAG-level abundance reconstruction

To assess the internal consistency of GORIN, we examined whether the reconstructed trajectories were consistent with the ASV copy-number structure encoded in each genome. A representative example is the MAG M601_h_023, which encodes three copies of ASV000103 and one copy of ASV000358 (see Additional file 1: Fig. S1b). Across the time series, the observed ASV counts maintained an approximately 3:1 ratio, matching the genome-encoded copy-number structure. The inferred MAG trajectory closely followed the temporal profile of the single-copy ASV, demonstrating internal consistency between the reconstructed MAG abundance and its constituent ASVs (see Additional file 1: Fig. S1c).

To assess internal consistency at the community scale, we further compared observed ASV time series with ASV trajectories reconstructed from the inferred MAG abundances for all genomes across both mice (see Additional file 1: Fig. S1d).

### Assessment of identifiability and cross-taxon ASV sharing

To evaluate the uniqueness of the inferred MAG abundances, we quantified the extent of rank deficiency in the ASV–MAG correspondence matrix. Connected components were classified as full-rank or rank-deficient, and the number of components, MAGs, and ASVs in each class was tabulated. We further calculated the proportion of total inferred MAG abundance attributable to MAGs in full-rank components.

We then examined the taxonomic ranks across which ASV sharing, the source of the abundance allocation handled by the network regularization, occurred among MAGs. Within rank-deficient components, we extracted MAG pairs that directly shared an identical ASV and compared their taxonomic assignments at the genus and family levels. At each rank, pairs in which both MAGs were assigned a taxon and the two taxa differed were retained for further examination.

Although all MAGs had passed the HMQ-MAG criteria, those belonging to these pairs were examined individually in greater detail. Specifically, we inspected completeness, contamination, GTDB-Tk classification, and the 16S rRNA gene sequences within each MAG together with their contig locations. MAGs showing evidence of chimeric assembly or binning contamination, and those with weak support for their taxonomic assignment, were designated as MAGs of questionable quality or assignment. When the problematic MAG within a group sharing an ASV could not be determined, all MAGs in that group were included in the assessment as unresolved assignments.

We calculated the proportion of total inferred MAG abundance accounted for by these MAGs separately for cross-genus and cross-family sharing, counting each MAG only once in each calculation. The abundance proportions for full-rank components and for MAGs of questionable quality or unresolved assignment were computed at each time point and summarized as the median and interquartile range across time points. The numbers of components, MAGs, and ASVs, together with the abundance proportions, are given in Additional file 1: Table S3.

### Functional gene composition estimation

To reconstruct time-resolved functional profiles, we combined genome-resolved functional annotations with the MAG abundance time series inferred by GORIN. For each annotation scheme, including KEGG Orthology and CAZyme families, we constructed a gene–MAG correspondence matrix *G*, where each entry *G*_*ij*_ represents the copy number of functional gene *i* encoded in MAG *j*.

Let ***x***(*t*) denote the inferred abundance vector of MAGs at time *t*. The functional composition at time *t* was computed as

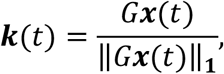

where ‖∙‖_**1**_ denotes the *L*_1_ norm, corresponding here to the sum of all elements of the nonnegative vector *G****x***(*t*). Thus, ***k***(*t*) represents the normalized functional composition at time t, while preserving genome-level linkage among features encoded within the same MAG.

### Short-read shotgun metagenomic sequencing

For short-read shotgun metagenomic sequencing, 24 fecal samples each from Mouse B and Mouse D (after vancomycin treatment) were selected. For each mouse, a 25-h window with few missing samples was selected from the hourly time series. Each selected window contained one missing time point, yielding 48 samples in total. Sequencing libraries were prepared using the ThruPLEX DNA-Seq Kit (Takara Bio USA, Inc.) following the manufacturer’s protocol. Sequencing was performed on the Illumina MiSeq i100 system to generate 150-bp paired-end reads.

### Construction and functional annotation of a short-read metagenomic gene catalog

Short-read shotgun metagenomic reads were processed sequentially using ParDRe [39] (v2.1.5) and fastp [40] (v0.23.4). Reads were then mapped against the Illumina PhiX (phiX174) reference and the mouse reference genome GRCm38.p6 (GCF_000001635.26) using minimap2[19] (v2.24). Reads identified by these mappings were excluded using SeqKit [41] (v2.3.0). Quality-controlled reads were assembled separately for each sample using metaSPAdes [42] (v4.3.0). Protein-coding genes were predicted from contigs ≥500 bp using Prodigal [31] (v2.6.3) in metagenomic mode (-p meta). Of the 6,960,613 predicted genes, 5,246,191 encoding proteins of at least 100 amino acids were retained. Their nucleotide sequences were clustered using CD-HIT-EST [43] (v4.8.1) at 95% sequence identity and an alignment coverage of at least 90% of the shorter sequence (-c 0.95 -aS 0.9), yielding a nonredundant catalog of 745,921 genes. KO annotations were assigned using KofamScan (v1.3.0) with the prokaryote.hal profile list and KO-specific adaptive score thresholds. A total of 201,326 genes (27.0%) were assigned to 3,659 KOs.

### Validation of inferred taxonomic and functional profiles

For taxonomic profiling, quality-controlled reads from each sample were subsampled to 3.5 million read pairs using seqtk [44](v1.5-r133) with a fixed random seed (-s42). Taxonomic relative abundances were estimated using MetaPhlAn4 [45] (v4.2.4) with the mpa_vJan25_CHOCOPhlAnSGB_202503 database.

MetaPhlAn profiles were converted to GTDB release R09-RS220 taxonomy using the MetaPhlAn utility sgb_to_gtdb_profile.py with the SGB-to-GTDB assignment table mpa_vJan25_CHOCOPhlAnSGB_202503_SGB2GTDB.tsv. To harmonize taxonomic assignments for this comparison, MAGs were reclassified against the same GTDB release using GTDB-Tk (v2.6.1). MetaPhlAn species-level abundances and GORIN-inferred MAG abundances were each aggregated at the genus level and matched by GTDB genus label. For Mouse B and Mouse D separately, the comparison used a genus set fixed across the 24 samples, comprising genera detected by either method and represented in the MetaPhlAn SGB-to-GTDB assignment table. Relative abundances from each method were retained without further renormalization; the MetaPhlAn unclassified fraction was excluded from the comparison but retained in its abundance denominator. Within this fixed genus set, nondetection in a sample was represented by zero. Spearman correlations were calculated on untransformed relative abundances across all genus–sample observations within each mouse, including observations with zero abundance in one or both profiles.

For functional profiling, reads from each sample were quantified against the nonredundant gene catalog using Salmon [46] (v2.7.0), using an index k-mer length of 31 and the quantification options -l A --validateMappings --seqBias --gcBias, with selective alignment and expectation– maximization-based abundance estimation. Gene-level TPM values estimated by Salmon were summed for each KO and normalized by the sum across all annotated KOs within each sample, yielding KO relative abundances that summed to one. The resulting shotgun-derived KO profiles were compared with those derived from GORIN-inferred MAG abundances and with those predicted by PICRUSt2 [47] (v2.6.3). PICRUSt2 predictions were generated from ASV representative sequences and abundance tables using the PICRUSt2-SC reference database and default KO prediction settings, including a maximum NSTI of 2.0 and normalization of ASV abundances by predicted 16S rRNA gene copy numbers. Comparisons were restricted to KOs present in both the PICRUSt2 and KofamScan reference databases.

Spearman correlations were calculated across all KO–sample observations within each mouse. In addition, Spearman correlations and Bray–Curtis dissimilarities were calculated separately for each sample. The GORIN-based and PICRUSt2-based KO profiles were compared using paired Wilcoxon signed-rank tests, with observations paired by sample within each mouse.

### Non-parametric period detection (NPPD)

Microbiome time-series data can contain irregular sampling, missing observations, non-sinusoidal waveforms, long-term drifts, and oscillatory dynamics whose period and phase vary over time. Global rhythmicity analyses summarize an entire time series by a single period and phase and therefore do not provide a locally varying phase trajectory. Because the downstream analyses in this study require phase estimates at each time point, we developed a Non-parametric Period Detection (NPPD) framework to reconstruct local phase trajectories without assuming a fixed waveform or stationarity (see Additional file 1: Fig. S3). NPPD quantifies local rhythmic transitions and reconstructs temporal phase trajectories without assuming a fixed periodicity or waveform, while accounting for statistical significance at each time point. The NPPD procedure consists of three steps: (1) computation of a local significance score based on Fisher’s exact test, (2) reconstruction of continuous phase trajectories from the score, and (3) identification of valid oscillatory cycles according to amplitude and duration criteria (see Additional file 1: Fig. S3).

The following sections describe each step of the method in detail.

### Local significance score for upward and downward transitions

To quantify local rhythmic transitions in each time series *x*(*t*), we defined a significance score based on Fisher’s exact test applied to the direction of short-term fluctuations relative to a 24-h trend (see Additional file 1: Fig. S3a).

For each series, a 24-h moving median was first computed to remove slow baseline drifts:

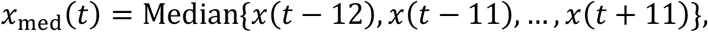

where the time resolution was one hour, and missing values or boundary indices were ignored. The detrended signal was then defined as:

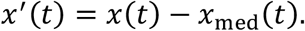

To determine whether the signal at time *t* represents a locally increasing or decreasing phase, we compared the signs of deviations in the preceding and following 12-h windows, using the local median of each window as the following threshold:

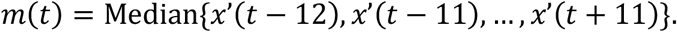

Then, the following quantities were defined to summarize the number of points above or below the threshold in each window:

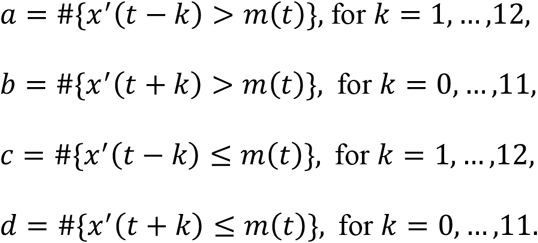

Fisher’s exact test was then performed on the contingency table 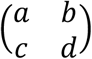 and the one-tailed *P*- value *P*_Fisher_(*t*) was used to define a signed local significance score:

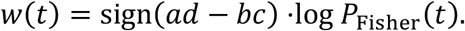

Positive *w*(*t*) values indicate significant upward transitions, whereas negative values indicate significant downward transitions. Alternating positive and negative peaks of *w*(*t*) approximately 12 h apart indicate a robust 24-h rhythmic oscillation in the original signal. This local, non-parametric detection method enables phase estimation even for noisy or irregularly sampled microbiome time series, without requiring global curve fitting or strict harmonic assumptions.

### Phase assignment from the local significance score

The temporal phase of each signal was determined from the zero-crossings and local extrema of the local significance score *w*(*t*) (see Additional file 1: Fig. S3b).

First, the *w*(*t*) time series was smoothed using a moving average filter to suppress high-frequency noise. The zero-crossing points were then identified according to changes in the sign of adjacent values. Transitions from positive to negative were classified as positive-to-negative crossings, and transitions from negative to positive as negative-to-positive crossings.

For each transition, the exact crossing time was estimated by linear interpolation between the two neighboring points. Consecutive zero-valued segments were treated as a single block; if the signs before and after the block differed, the block was assigned to the corresponding crossing type.

As the signal necessarily starts and ends at zero by definition, the endpoints were explicitly assigned to a crossing type based on the sign of the adjacent segment.

Next, within each interval bounded by two consecutive zero-crossing points, we determined either the local maximum or minimum of *w*(*t*), depending on the sign of the interval.

Specifically, for intervals between a positive-to-negative and a subsequent negative-to-positive crossing, the minimum value of *w*(*t*) was identified as a local trough, denoted *w*_trough_ ; whereas for intervals between a negative-to-positive and a subsequent positive-to-negative crossing, the maximum value of *w*(*t*) was identified as a local peak, denoted *w*_peak_. These extrema served as anchor points that represent the timing of the upward and downward phases of the 24-h oscillation. The temporal sequence of peaks and troughs thus defines the phase progression of the periodic behavior for each signal.

The temporal sequence of zero-crossings and local extrema of *w*(*t*) was mapped onto a continuously increasing unwrapped phase *ϕ*_*u*_(*t*). Each oscillatory cycle was divided into four consecutive segments defined by the temporal order of the local extrema:

1. a positive-to-negative crossing (*ϕ*_*u*_ = 0),
2. a local trough (*ϕ*_*u*_ = *π*/2),
3. a negative-to-positive crossing (*ϕ*_*u*_ = *π*), and
4. a local peak (*ϕ*_*u*_ = 3*π*/2).

Within each segment, the phase increased linearly with time between the two bounding reference points. When the next cycle began, 2*π* was cumulatively added to maintain monotonicity across cycles, yielding an unwrapped continuous phase trajectory *ϕ*_*u*_ (*t*).

The instantaneous phase *θ*(*t*) was then obtained by wrapping *ϕ*_*u*_ (*t*) into the principal interval [0,2π) as

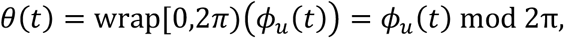

where *θ* = 0 and *θ* = 2*π* denote the same phase point on the unit circle.

For pairwise phase-difference calculations, the wrapped difference was projected onto the centered interval (−π, π] as

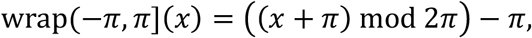

ensuring continuity near the ±*π* boundary. A complete oscillatory cycle was defined as the temporal sequence of a positive-to-negative zero-crossing, trough, negative-to-positive zero-crossing, and peak, returning to the next positive-to-negative zero-crossing. This sequence corresponds to a continuous phase advance of 2*π*.

### Detection of valid cycles

After identifying peak and trough anchor points, we divided each time series into individual oscillatory cycles. Cycle boundaries were defined by the phase corresponding to the troughs (*θ* = *π*), such that each cycle spans from one trough to the next (−*π* to *π*), corresponding to a one daily cycle. Each candidate cycle was evaluated according to the following criteria, and only those satisfying all conditions were retained as valid cycles (see Additional file 1: Fig. S3c):

1. Amplitude: The mean of the absolute values of the local peak and trough exceeded 1.0, i.e.,

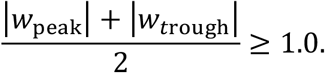
2. Duration of the cycle: The time interval between consecutive troughs was between 16 and 32 h.
3. Mean abundance (for MAG abundance data): The average relative abundance of MAG during the cycle exceeded 0.01% of the total abundance.

These thresholds ensure that only robust, biologically meaningful oscillations are analyzed, excluding weak or irregular fluctuations. The resulting set of valid cycles was subsequently used for downstream phase alignment and population-level analyses. An implementation of the NPPD framework used in this study is publicly available on GitHub at https://github.com/rie-maskawa/Non-parametric-Period-Detection.

### Phase benchmark against global rhythmicity analyses

To benchmark local phase estimation, we compared NPPD with JTK_CYCLE [48] and the Lomb–Scargle periodogram [49,50]. JTK_CYCLE was run using the MetaCycle implementation [51], and both methods searched periods of 16–32 h, corresponding to the range in which NPPD validates cycles. JTK_CYCLE provided the fitted period and lag, whereas for Lomb–Scargle the phase was obtained from a cosinor fit at the periodogram peak. For the synthetic benchmark, we generated 200 time series of 336 hourly points. In each replicate, the oscillation period changed linearly over 14 days from a value drawn between 16 and 20 h at the beginning of the record to a value drawn independently between 28 and 32 h at the end, such that the true instantaneous phase was known at every time point and no single period described the complete series. Within each cycle, log relative abundance followed a fixed asymmetric waveform, rising over the first third of the cycle and declining over the remaining two thirds, with an amplitude of 0.45 in natural-log units around a baseline relative abundance of 0.005. Correlated noise was added in log space as an AR(1) process with ρ = 0.8 and scale 0.35. Each hourly point was additionally assigned an outlier with probability 0.01 and multiplied, with equal probability, by a factor drawn uniformly from 3–8 or 0.05–0.3. Counts were then sampled from a negative-binomial distribution parameterized as Var(*Y*) = µ + ϕµ^2^, with dispersion ϕ = 0.08 and a library size of 30,000 reads, and converted to relative abundances. Oscillation amplitude and noise level were held constant throughout each simulation, so that only the local period changed systematically over time.

Circular phase errors were converted to hours using the true local period at each time point. The first and last 12 h of each record were excluded because the centered rolling window used by NPPD was not fully defined at the boundaries. Phase-estimation accuracy was summarized as the fraction of pooled time points with an absolute error no greater than a tolerance τ.

### Hierarchical clustering based on phase difference

To identify groups of taxa exhibiting similar temporal phase relationships, we performed hierarchical clustering based on the mean phase difference between all pairs of oscillatory time series. For each pair of entities *i* and *j* with instantaneous phases *θ*_*i*_(*t*) and *θ*_*i*_(*t*), the phase difference was defined as

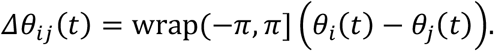

We then defined the pairwise distance between entities as the temporal average of the absolute circular phase difference,

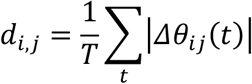

This metric quantifies the mean angular separation between two oscillatory signals, yielding *d*_*i,j*_ = 0 for perfectly synchronized oscillations and *d*_*i,j*_ = *π* for oscillations in anti-phase. The resulting distance matrix was used to perform hierarchical clustering of taxa.

To minimize edge effects caused by the start and end of the observation window, the first and last 12 h of each time series were excluded from the calculation of the temporal average. The pairwise distance matrix was subjected to hierarchical clustering using the average linkage method, and phase-synchronized clusters were defined from the resulting dendrogram structure.

### Functional enrichment analysis between phase clusters

To evaluate functional biases between the two phase-defined microbial clusters, we used the gene–MAG copy number matrix 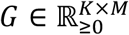 where K and M denote the total numbers of genes (e.g., KEGG Orthologs or CAZy families) and MAGs, respectively, and each element *G*_*ij*_ represents the copy number of gene *i* in MAG *j*. Let *C*_Light_ and *C*_Dark_ denote the sets of MAGs assigned to the Light and Dark clusters, respectively, according to phase-based classification. For each gene *i*, we defined two subsets of copy numbers:

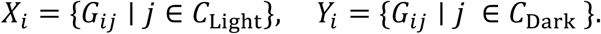

Genes with all-zero entries in both, were excluded from further analysis. Differences between groups were assessed using the Mann–Whitney *U* test (two-sided, asymptotic approximation). Effect size for gene *i* was quantified by Cliff’s delta,

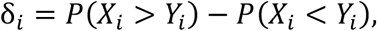

which measures the probability difference that values from the light group exceed those from the dark group. Ties contribute zero to δ_*i*_, and positive and negative values indicate higher copy numbers in the Light and Dark clusters, respectively. Multiple testing correction was performed across genes using the Benjamini–Hochberg false discovery rate (FDR) procedure. Genes were defined as significantly enriched if they met both criteria: FDR-adjusted *P* < 0.05 and an absolute Cliff’s delta (|*δ*|) ≥ 0.25.

To evaluate enrichment at higher functional levels, gene-level results were aggregated by KEGG pathways and modules. For each pathway/module o, a 2×2 contingency table was constructed comparing the number of significant genes within and outside o, and significance was assessed using Fisher’s exact test (one-sided, testing for over-representation). A pathway or module was considered enriched when *P* < 0.05, the Haldane–Anscombe-corrected odds ratio exceeded 3 for pathways or 5 for modules, and at least two member genes were significant. The analysis was performed separately for Mouse A and Mouse B, and only pathways and modules enriched in both mice were reported.

To assess whether the functional differences between the Light and Dark clusters were consistent across individual phyla, we repeated the analysis separately within each phylum represented in both clusters. MAGs from the six mice were pooled because individual mice often contained too few MAGs from a given phylum in one of the two clusters for within-phylum comparison. Phyla containing at least two Light and two Dark MAGs were included (*Bacteroidota, Bacillota_A*, and *Bacillota*). Within each phylum, gene copy numbers were compared between Light and Dark MAGs using the same Mann–Whitney U test and Cliff’s delta as in the cluster-level analysis, with Benjamini–Hochberg correction applied across the genes tested within that phylum. KEGG pathway and module enrichment was then assessed from the resulting cluster-associated genes using the same one-sided Fisher’s exact test and set-level thresholds as in the main analysis, with the background restricted to the genes tested within that phylum. CAZyme families were similarly compared between Light and Dark MAGs within each phylum using the same gene-family-level procedure.

### Circular correlation of peak phases across mice

To quantify the conservation of peak-phase relationships across mice, we calculated the Jammalamadaka–Sengupta circular correlation coefficient, ρ_*c*_ [52], for features shared between each pair of mice. For paired peak phases θ_*i*_ and ϕ_*i*_,

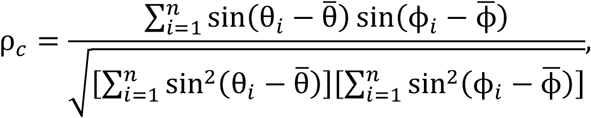

where 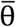 and 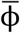 denote the circular mean peak phases in the two mice. The coefficient ranges from -1 to 1, with values approaching 1 indicating a strong positive circular association, values near 0 indicating little circular association, and values approaching -1 indicating a strong negative circular association. Because ρ_*c*_ is invariant to a uniform rotation of either set of phases, it quantifies the conservation of relative peak-phase relationships even when the overall peak timing is shifted between mice. This property was particularly suited to our analysis, in which we aimed to assess the preservation of relative temporal organization rather than absolute peak-time agreement.

Statistical significance was evaluated using the asymptotic test for the null hypothesis *H*_0_: ρ_*c*_ = 0 described by Rao and SenGupta [52]. Defining

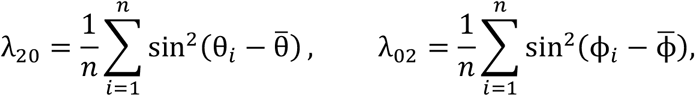

and

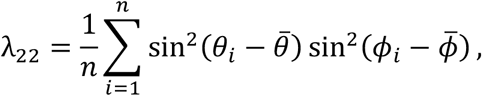

the test statistic was calculated as

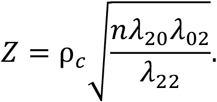

Under *H*_0_, *Z* is asymptotically standard normally distributed, and two-sided *P*-values were calculated accordingly.

## Results

### Automated fecal collection enables high-frequency microbiome time-series sampling

We developed an automated system that enables the long-term and high-frequency collection of mouse feces without manual intervention. The device consists of three main components—a metabolic cage, a fraction collector, and a control unit (Fig. 1a; see Additional file 1: Supplementary Notes). The metabolic cage separates feces and urine and directs them to distinct tubes positioned below the funnel. Each fecal tube is prefilled with RNA*later*™ (Fig. 1b), a reagent known to preserve the microbiome composition [53], and the collector rotates beneath the drop point at user-defined intervals ranging from minutes to hours, with proof-of-concept experiments demonstrating stable sampling at 5- and 10-min resolution (see Additional file 1: Fig. S4). After each collection run, a stainless-steel ball is released from a spiral feeder via a swing arm onto the tube, preventing contamination from subsequently falling material and minimizing RNA*later*™ or urine evaporation (Fig. 1c).

**Fig. 1.**
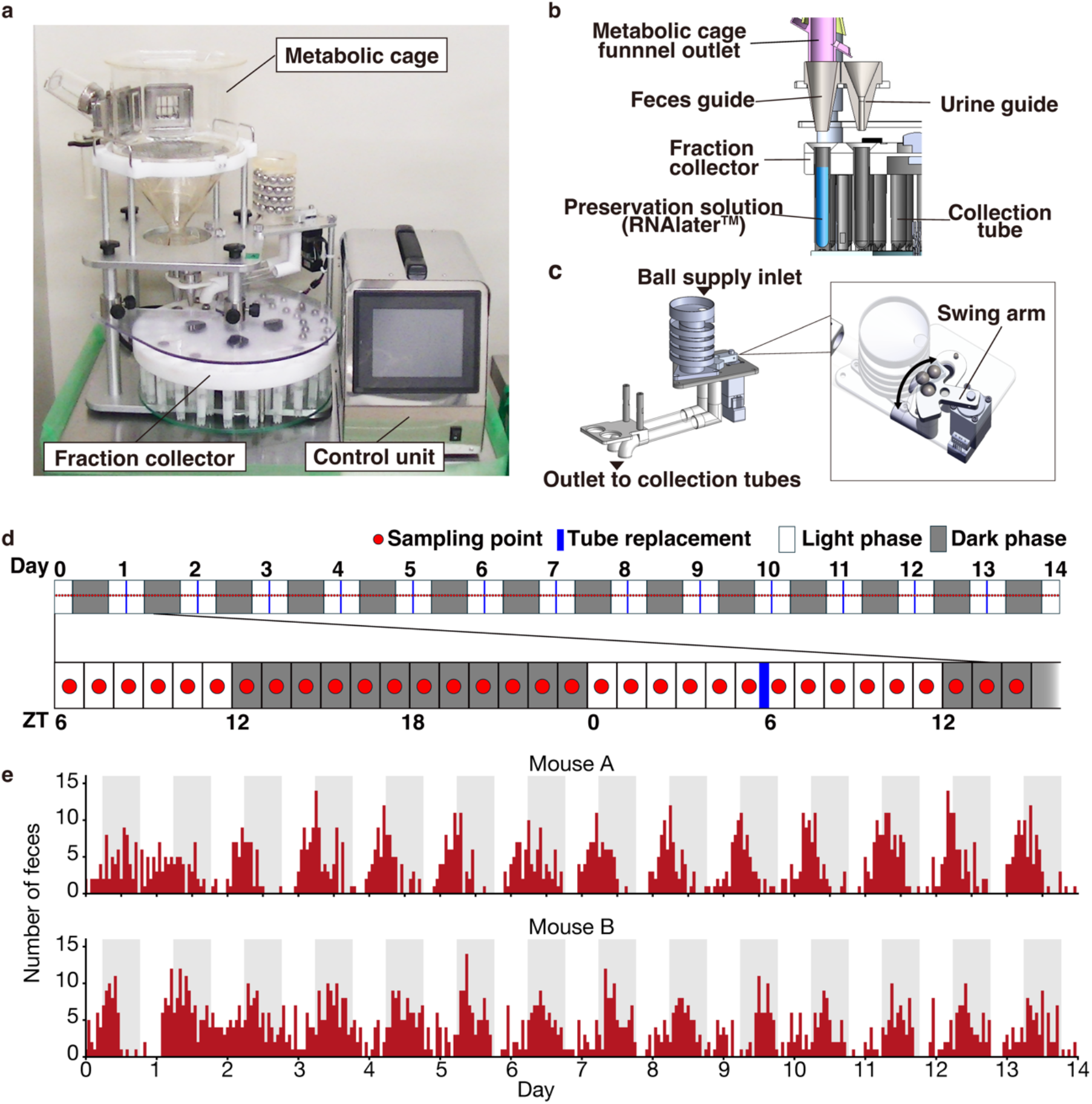
Development of an automated fecal collection system and high-frequency microbiome time-series sampling. (a) Photograph of the automated mouse fecal collection system, which consists of a metabolic cage, fraction collector, and control unit. (b) Schematic illustration of the structures located beneath the metabolic-cage funnel outlet. Feces and urine are separated by dedicated guiding funnels, directing fecal pellets into collection tubes pre-filled with the preservation solution (RNA*later*™) and urine into independent tubes within the fraction collector. (c) Stainless-steel ball supply mechanism and the underlying swing-arm structure. Balls are stacked in the supply inlet and sequentially delivered through the swing-arm mechanism to the outlet that leads to the collection tubes. (d) Sampling schedule for 14 consecutive days. Each red circle indicates an hourly sampling point, and blue bars indicate the tube replacement timing. Light and dark phases correspond to the 12:12-h light/dark cycle. (e) Hourly counts of fecal pellets collected over 14 days. Bar plots show the number of pellets recovered each hour for Mouse A (top) and Mouse B (bottom). Gray shading indicates the dark phase.

Using this system, we continuously collected feces from two mice (Mouse A and Mouse B) at 1-h intervals over 14 days (Fig. 1d). The collection process was fully automated throughout each sampling cycle, requiring only brief daily maintenance for tube replacement and funnel cleaning. Each hour, naturally excreted fecal pellets (0–14 pellets per mouse per hour) were automatically collected (Fig. 1e). Because fecal sampling was performed directly within the metabolic cage, collection proceeded without removing the animals, simplifying routine operations throughout the experiment. The high-frequency fecal output trace showed pronounced hour-to-hour fluctuations in natural defecation and a clear tendency for more frequent pellet excretion during the dark phase. This dataset had substantially higher temporal resolution and longer continuity than previous data [4], thus successfully capturing intrinsic defecation dynamics unobservable with conventional, sparsely collected data.

We compared automated and manual fecal collection in four additional mice at two sampling times, yielding eight matched pairs. Genus-level relative abundances were highly correlated between workflows (median Spearman’s ρ = 0.946), and the median ASV-level Bray–Curtis distance between workflows was smaller than those for within- and between-mouse comparisons. In hierarchical clustering, each automatically collected sample clustered first with its matched manually collected sample in all eight pairs (see Additional file 1: Fig. S5 and Supplementary Notes).

### Reconstructing genome- and function-level time series from full-length 16S rRNA profiles

Fecal pellets were processed for full-length 16S rRNA gene and long read metagenomic sequencing. Using long-read assemblies as a reference, each genome was matched to its corresponding ASVs inferred from the 16S rRNA gene sequence reads, generating the copy-number matrix C as a direct ASV–MAG association (Fig. 2a and see Additional file 1: Fig. S6). Using this copy-number matrix, we reconstructed genome-resolved abundance trajectories from ASV counts with Genome abundance Optimization from 16S rRNA-Informed Networks (GORIN). In GORIN, the relationship between observed ASV counts and MAG abundances was modeled as

**Fig. 2.**
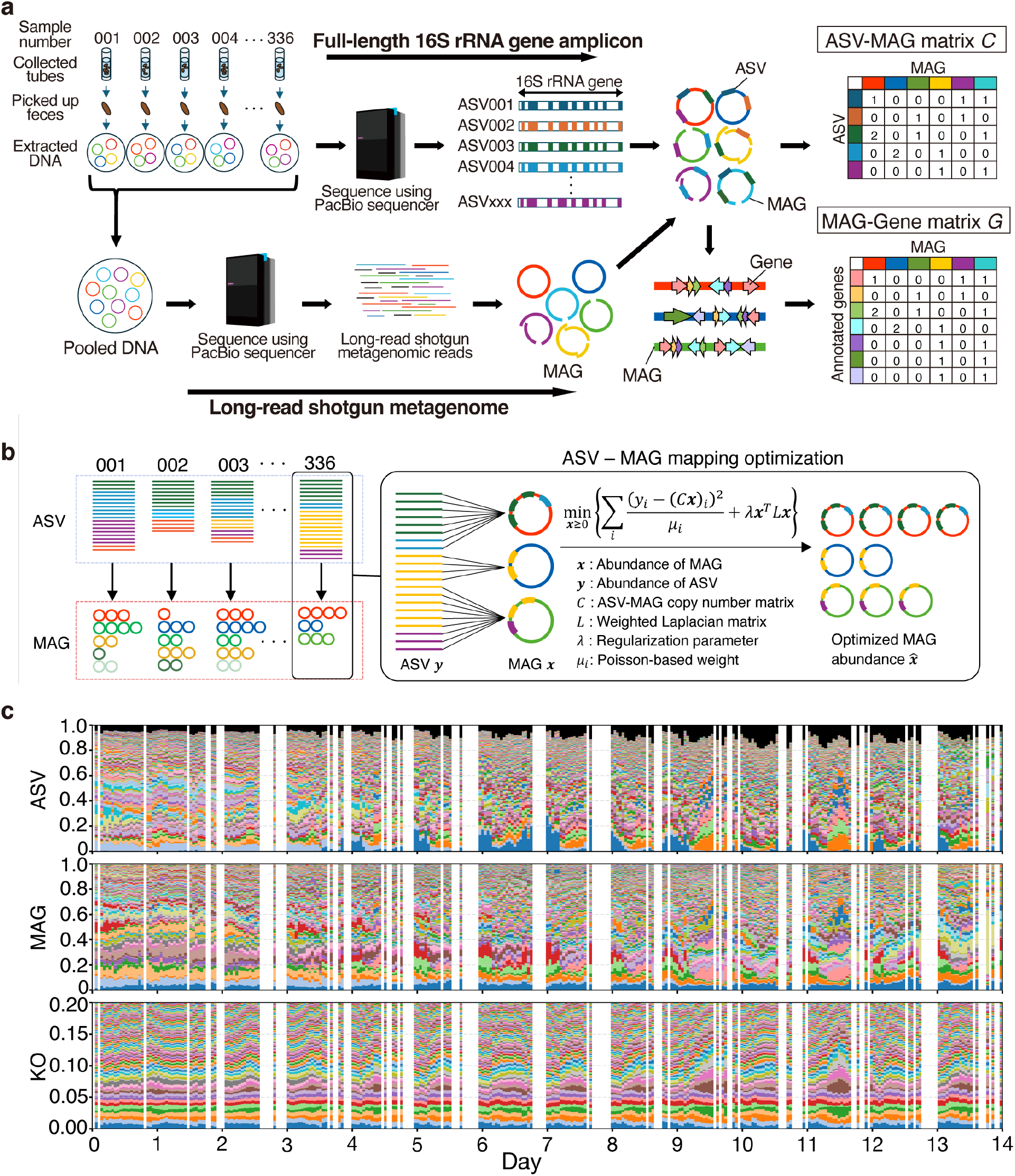
Estimation framework for MAG abundance and functional gene dynamics from full-length 16S rRNA profiles using GORIN. (a) Workflow for the integrated microbiome analysis. Fecal samples collected at each time point were subjected to full-length 16S rRNA gene amplicon sequencing, generating ASVs, whereas pooled time-series samples were used for long-read metagenomic sequencing to reconstruct MAGs. The ASV–MAG matrix was constructed based on exact sequence matches between ASVs and MAGs and the gene–MAG matrix was generated to link assembled genomes with annotated functional genes. (b) Overview of GORIN, the optimization framework for estimating MAG abundances from observed ASV counts, using the ASV–MAG copy number matrix. (c) Overview of reconstructed dynamics across hierarchical levels for Mouse A: relative abundances of ASVs, MAGs, and KEGG orthologs (KOs). Components in each stacked plot are ordered by mean abundance, and the KO panel is shown with an expanded vertical scale. Black areas in the ASV panel represent ASV reads not assigned to any MAG, with 91.8% of total ASV reads successfully associated with reconstructed MAGs.

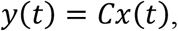

where *y*(*t*) and *x*(*t*) denote the ASV count vector and corresponding MAG abundance vector at time *t*, and *x*(*t*) was estimated by constrained optimization (Fig. 2b) (details in Methods). The ASV trajectories reconstructed from the inferred MAG abundances were consistent with the observed ASV dynamics, supporting the internal consistency of GORIN (see Additional file 1: Fig. S1d).

We further annotated genes in each MAG to construct a MAG–gene matrix (G) and used it with the estimated MAG abundances to compute time-resolved functional profiles. Fig. 2c summarizes the reconstructed time series across hierarchical levels—ASV, MAG, and KEGG Orthologs [32] (KOs)—ordered by mean abundance. The corresponding results for the other five mice are shown in Additional file 1: Fig. S7. To independently assess the inferred taxonomic and functional profiles, we generated short-read shotgun metagenomic data for 24 samples each from Mouse B and Mouse D. Genus-level abundances aggregated from the inferred MAG profiles correlated with MetaPhlAn estimates, with Spearman correlations of 0.73 and 0.75 across all genus–sample observations for Mouse B and Mouse D, respectively (see Additional file 1: Fig. S2a). At the functional level, inferred KO relative abundances showed correlations of 0.94 and 0.92 with the shotgun-derived profiles across all KO–sample observations, compared with 0.68 and 0.69 for PICRUSt2 for Mouse B and Mouse D, respectively (see Additional file 1: Fig. S2b). Sample-wise comparisons also showed higher Spearman correlations and lower Bray–Curtis distances to the shotgun-derived KO profiles for the GORIN-based profiles than for PICRUSt2 (see Additional file 1: Fig. S2c–d). These results support the inferred genus-level and functional compositions. Together, GORIN enabled the reconstruction of genome- and function-level time series from ASV profiles for subsequent analyses of the temporal organization of the gut microbiome.

### Multiscale taxonomic and functional composition temporal dynamics

To characterize the compositional dynamics across hierarchical levels, we performed a principal coordinates analysis (PCoA) based on Bray–Curtis distances of the metagenomic time series at both the MAG and KO levels (Fig. 3a–b and see Additional file 1: Fig. S8a–b). The MAG level profiles exhibited both a slow multi-day drift (PCo1, ∼34% variance), and a 24-h cycle (PCo2, ∼17%) (Fig. 3a and see Additional file 1: Fig. S8a), whereas, at the KO level, PCo1 (∼53% variance) was dominated by the diurnal cycle and showed little evidence of multi-day drift, suggesting that functional profiles fluctuate within a much narrower range (Fig. 3b and see Additional file 1: Fig. S8b). Temporal autocorrelation analysis confirmed clear daily periodicity in both datasets, but multi-day divergence accumulated only at the taxonomic level, indicating substantially greater stability of functional composition (Fig. 3c and see Additional file 1: Fig. S8c). Consistent with this community-level pattern, the overall variability of individual KOs was significantly lower than that of individual MAGs across all time scales (see Additional file 1: Fig. S8d–e).

**Fig. 3.**
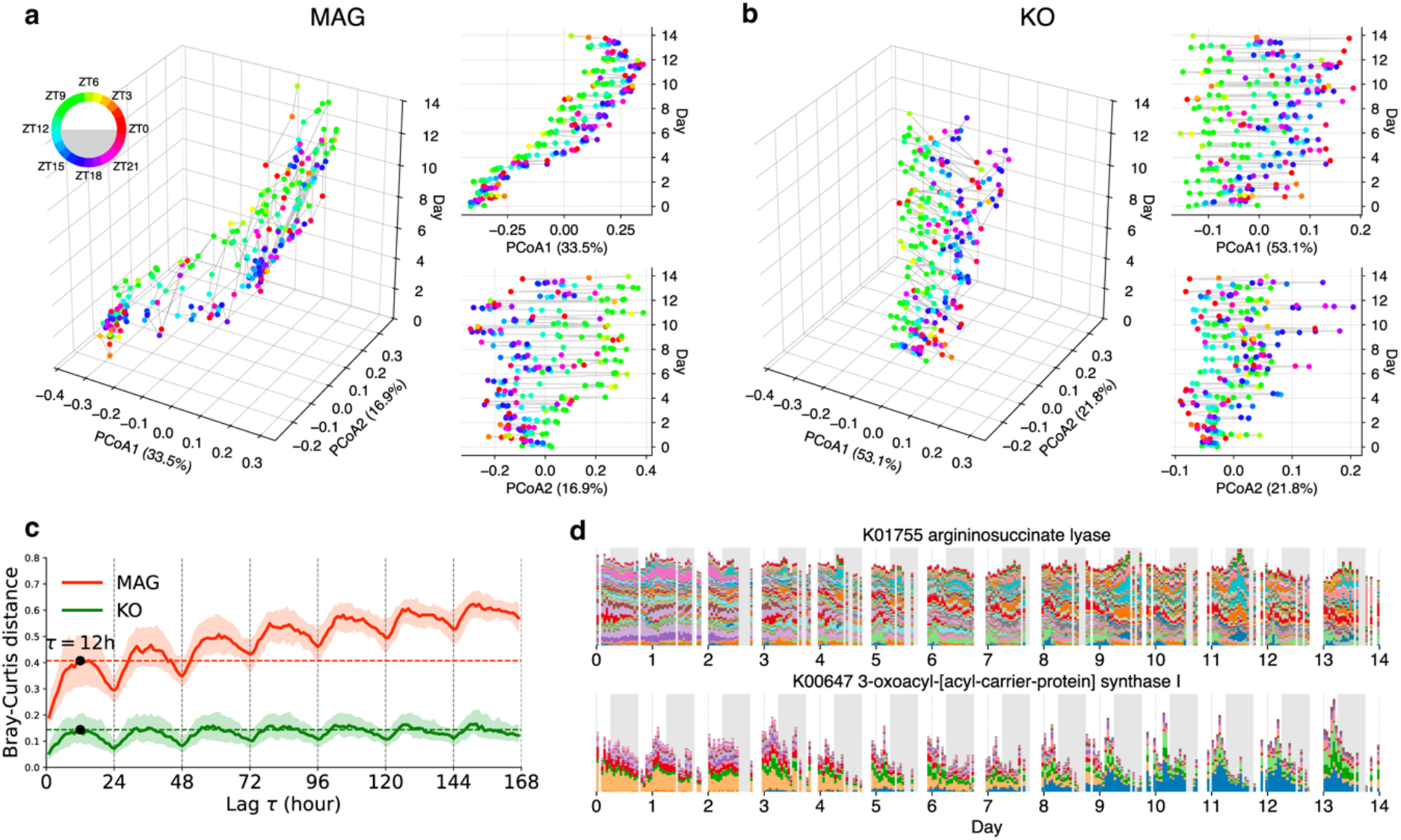
Multiscale temporal dynamics of taxonomic and functional compositions. (a–b) Principal coordinates analysis (PCoA) based on Bray–Curtis distances of metagenomic time-series data at the MAG (a) and KO (b) levels. Each point represents a sample colored by the sampling time (ZT). Left: three-dimensional plots with PCo1–PCo2 on the horizontal axes and time (day) on the vertical axis. Right: temporal trajectories of principal coordinates. (c) Temporal Bray–Curtis dissimilarity as a function of time lag (τ), representing the temporal autocorrelation structure of the community composition at the MAG (orange) and KO (green) levels. Solid lines indicate the median values across all time windows, and shaded areas represent the interquartile range. (d) Examples of KO-level time series reconstructed from MAG abundances. Each colored band represents the contribution of individual MAGs to the total abundance of a given KO. The top panel shows K01755 as an example of a function supported by multiple MAGs, and the bottom panel shows K00647 as a function associated with fewer major contributors.

The observed functional gene dynamic stability could result from distinct underlying mechanisms, including statistical averaging due to functional redundancy among diverse taxa, known as a portfolio effect [54] (see Additional file 1: Fig. S8f). For example, the arginine biosynthesis enzyme K01755 (*argH*) represents highly redundant functions supported by numerous genomes (220 MAGs), where simultaneous contributions from multiple taxa collectively maintained a smooth temporal trajectory (Fig. 3d (top)). In contrast, the function of fatty-acid synthase K00647 (*fabB*) is supported by a much smaller set of contributors (15 MAGs). Although the identities of the contributing MAGs changed markedly over time, their complementary fluctuations preserved a stable functional profile (Fig. 3d (bottom)). These observations suggest that functional stability in the gut microbiome can emerge through both averaging among redundant contributors and compensatory dynamics among specialized taxa.

### Phase analysis revealed coordinated circadian organization of the gut microbiome

To resolve the circadian structure in microbiome dynamics, we introduced a phase representation, mapping each time point to an angular position θ on the unit circle (Fig. 4a). Although several approaches have been proposed to infer periodic structure directly from time-series data [48], they assign a single period and phase to an entire record, so a trajectory whose period drifts, or whose oscillation occupies only part of the record, is not represented by their output (see Additional file 1: Fig. S9). To overcome this, we developed a new data-driven phase-analysis framework, NPPD (see Additional file 1: Fig. S3). NPPD reconstructs oscillatory cycles and assigns a phase at every time point along those cycles, without assuming a fixed waveform or stationarity (Fig. 4a). Consistent with this purpose, benchmarking against two representative global approaches, JTK_CYCLE and Lomb–Scargle, showed that NPPD more closely tracked the true phase when the oscillation period changed over time (see Additional file 1: Fig. S10). NPPD includes a statistical procedure that evaluates the validity of each reconstructed 24-h cycle, allowing us to identify reliably oscillating components. The distribution of valid cycles per MAG reveals that many members of the community display daily rhythmicity, ranging from intermittent to sustained oscillations (see Additional file 1: Fig. S11a). For Mouse A and Mouse B, we defined core oscillatory MAGs as those with ≥ 7 valid cycles, that is, oscillating on at least half of the recorded days. This set comprised 118 of 329 MAGs in Mouse A and 181 of 463 MAGs in Mouse B, accounting for 71% and 60%, respectively, of the MAG-level relative abundance averaged across the observation period. All subsequent phase-based analyses were performed exclusively on this core set.

**Fig. 4.**
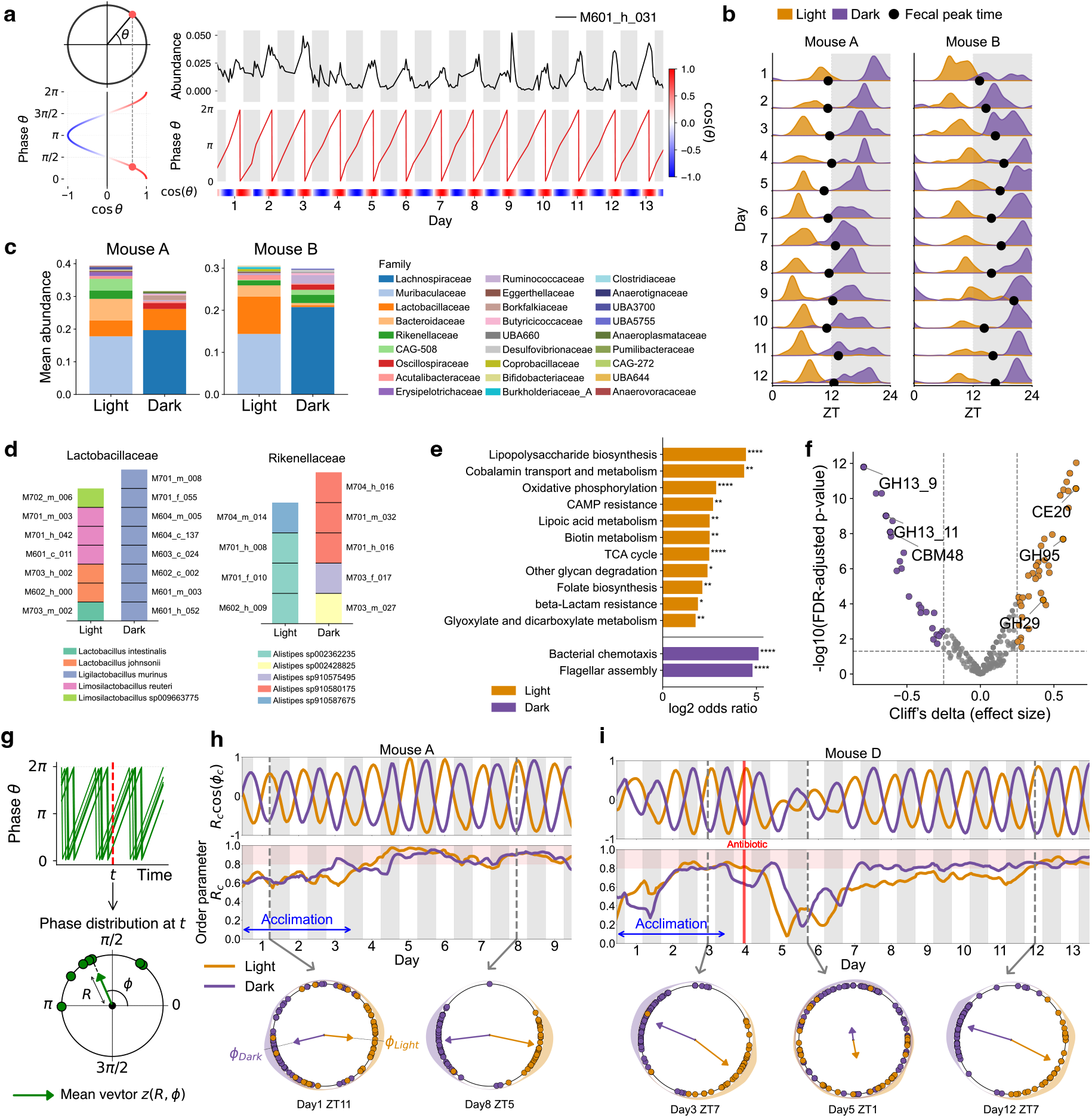
Phase-based characterization of circadian organization of gut microbial communities. (a) Schematic of the phase representation: each time point is mapped to an angular position θ on the unit circle (Left). Example of phase reconstruction for a representative MAG (M601_h_031), showing the relative abundance, estimated phase θ, and cos(θ) (Right). (b) Distribution of peak times of oscillatory MAGs in mice A and B, colored by cluster assignment (Light, orange; Dark, purple). Peak time corresponds to phase θ = 0. Black dots denote fecal excretion peaks. (c) Mean family-level composition of the Light and Dark clusters. (d) Species composition of the families *Lactobacillaceae* (left) and *Rikenellaceae* (right) in the Light and Dark clusters. Stacked bars indicate the MAGs belonging to each family detected in the clusters, including genomes derived from both mouse A and mouse B. (e) KEGG pathways significantly enriched in each cluster (one-sided Fisher’s exact test with the set-level criteria described in Methods; enriched in both mice). (f) Differential CAZy enrichment between clusters. Each point represents a CAZy family (x-axis: Cliff’s delta; y-axis: FDR-adjusted *P*- value). (g) Schematic illustration of the mean phase vector, *z* = *Re*^*iϕ*^ . (h–i) Temporal evolution of phase alignment for the Light (orange) and Dark (purple) clusters in two representative mice: (h) Mouse A, and (i) Mouse D in which a single dose of antibiotics was administered on day 4. The top panels show the real part of the mean phase vector, *R*_*c*_ cos(ϕ_*c*_), which represents the directional projection of the cluster mean phase on the unit circle. The second panels show the time course of the order parameter *R*_*c*_, which quantifies the degree of phase coherence among genomes within each cluster. The bottom circular plots display the phase distribution of individual genomes at representative time points (indicated by grey dashed lines); arrows denote the mean phase vector, whose length corresponds to *R*_*c*_.

Pairwise phase differences (Δθ) among oscillatory MAGs exhibited a clear bimodal distribution, indicating the presence of two dominant phase relationships within the community (see Additional file 1: Fig. S11b). Consistent with this structure, hierarchical clustering based on pairwise phase distances separated the community into two major phase groups. The peak-time distribution of oscillatory MAGs showed a corresponding bimodal pattern, with peaks concentrated in the light (ZT 0–12) and dark (ZT 12–24) phases (Fig. 4b). Based on these peak times, the two groups were designated the Light and Dark clusters. Despite day-to-day and individual variation in absolute peak timing, fecal excretion peaks were consistently positioned between the two cluster distributions, close to the midpoint of the Light and Dark peak times. The same bimodal structure was observed when absolute abundances were analyzed, confirming that the pattern was not driven by compositional effects (see Additional file 1: Fig. S12).

### Taxonomic and functional differentiation between Light- and Dark-phase clusters

High-abundance MAGs commonly exhibited distinct compositions between Light and Dark clusters. The Light cluster was dominated by *Bacteroidota*, including *Muribaculaceae* and *Bacteroidaceae*, whereas the Dark cluster was dominated by *Bacillota*, including *Lachnospiraceae* and *Oscillospiraceae* (Fig. 4c), consistent with previous reports [4]. At the family level, *Lactobacillaceae* and *Rikenellaceae* were abundant in both clusters. However, species within these families were completely segregated between Light and Dark clusters (Fig. 4d). These patterns were also observed in Mouse C, Mouse D, Mouse E, and Mouse F (see Additional file 1: Fig. S13a–b). This clear species-level separation was revealed through our genome-level community structure reconstruction.

To further identify functional features of each cluster, we identified genes that were commonly enriched among mice. In the Light cluster, genes involved in lipopolysaccharide biosynthesis, a hallmark of gram-negative bacteria, endogenous cationic antimicrobial peptide resistance and vitamin biosynthesis were enriched (Fig. 4e and see Additional file 1: Fig. S13c). Consistent with these features, CAZyme [34] families, involved in mucopolysaccharide degradation, such as CE20, GH95, and GH29, were specifically enriched in the Light cluster (Fig. 4f).

In contrast, the Dark cluster was enriched in genes related to bacterial chemotaxis, flagellar assembly, and carbon storage metabolism, including pathways for trehalose and starch biosynthesis (Fig. 4e and see Additional file 1: Fig. S13c). Consistently, several CAZyme families involved in carbohydrate accumulation and storage, such as GH13_9, GH13_11, and CBM48, were specifically enriched in the Dark cluster (Fig. 4f).

To distinguish phase-associated functional differences from lineage-associated differences in gene content, we further assessed whether the functional differences observed between the Light and Dark clusters were consistent within individual phyla represented in both clusters (Additional file 1: Fig. S14). Several features enriched in the original cluster-level comparison, including LPS-related and transport functions, were no longer supported within phyla, indicating that part of the original signal reflected lineage-associated differences in gene content. In contrast, many functional differences remained detectable within individual phyla. For example, bacterial chemotaxis and flagellar assembly remained strongly enriched in the Dark cluster within *Bacillota_A*, together with multiple carbohydrate-related functions. Moreover, several Dark-directed features were significant in both *Bacillota_A* and *Bacillota*, including trehalose biosynthesis and starch- and glycogen-related CAZyme families. Some Light-directed features, such as GH29, were significant within *Bacteroidota* and showed the same direction in another phylum. Together, these recurrent patterns indicate a phase-associated functional bias shared across taxonomically distinct lineages.

### Emergence of synchronized phase dynamics

To quantify the degree of phase alignment within each cluster, we introduced the mean phase vector:

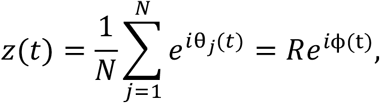

where *θ*_*j*_(*t*) denotes the instantaneous phase of the *j*-th MAG at time *t* (Fig. 4g). For each cluster *c* (Light or Dark), we computed the cluster-specific mean phase vector *z*_*c*_(*t*) . Its magnitude, *R*_*c*_ (*t*) = |*z*_*c*_ (*t*)|, represents the phase order parameter, quantifying the coherence among members of cluster c, whereas *ϕ*_*c*_ (*t*) denotes the corresponding mean phase [55].

This synchronization analysis of the two genome clusters revealed a coordinated transition from initially dispersed to highly synchronized phases following cage transfer. Immediately after the transfer, phase coherence was low and the phases of individual MAGs were widely dispersed, but around day 4, coinciding with host acclimation, phase coherence began to increase steadily, and the relative phases narrowed markedly (Fig. 4h). Similarly, in Mouse D, which received a single oral dose of vancomycin (day 4 at 15:00), phase synchrony was markedly disrupted after one day of treatment and gradually recovered over the following 5 days (Fig. 4i). None of these phase-based results depends on the valid-cycle criterion: relaxing it stepwise to any MAG with at least one valid cycle preserves the two-cluster structure, the antiphase relationship between the clusters, their taxonomic and functional contrast and the time course of the order parameter, and lowers only its absolute level (Additional file 1: Fig. S15).

### Diel succession of microbial taxa and functions

Temporal phase analysis of hourly-resolved microbial taxa and gene functions revealed not only the separation of the microbiome into light- and dark-phase groups but also a reproducible temporal succession within each phase (Fig. 5a–b and see Additional file 1: Fig. S16a). Ordering microbial taxa and gene functions by peak phase demonstrated a gradual progression from early to late within each circadian phase that was conserved across mice.

**Fig. 5.**
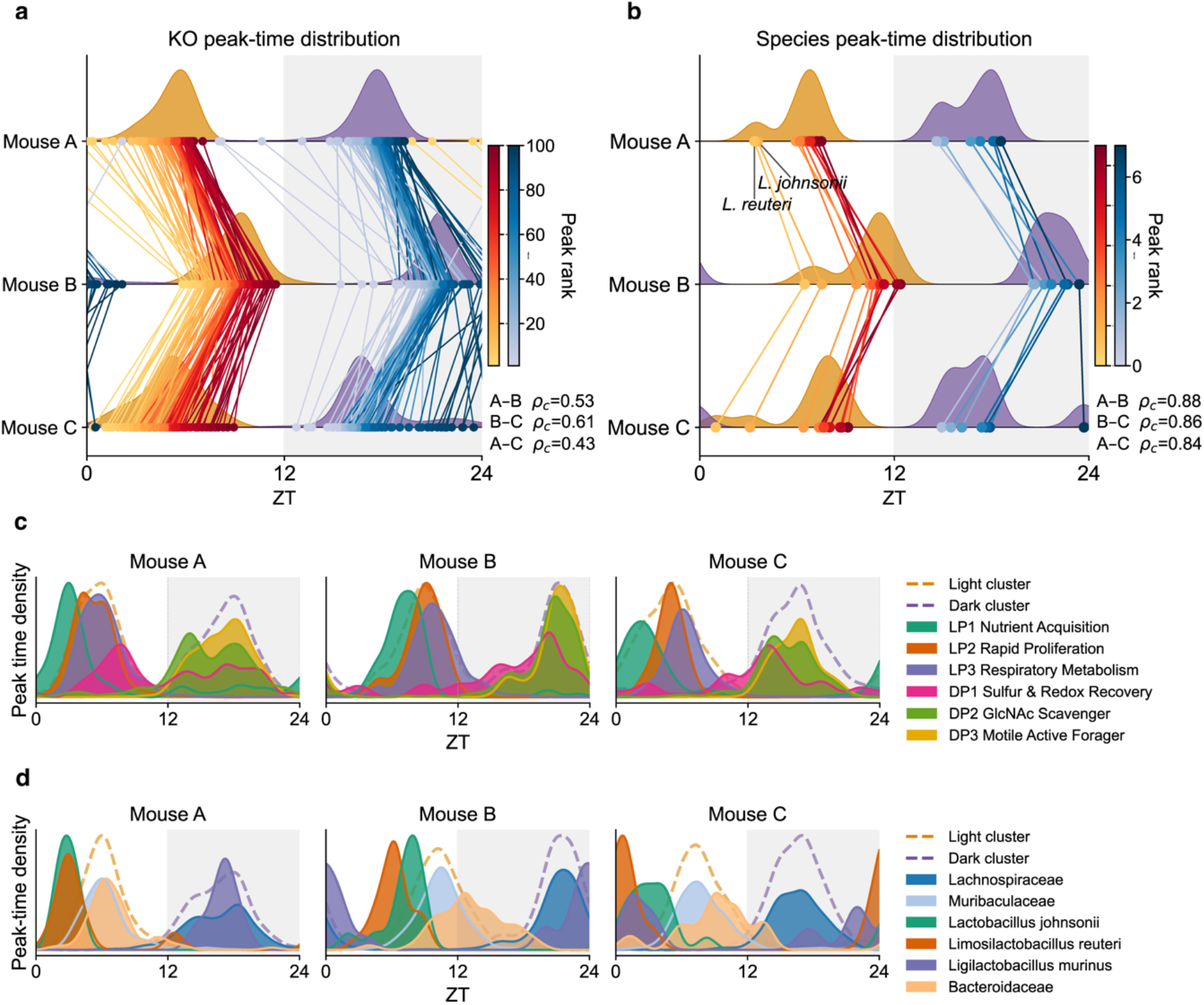
Coordinated diel organization of microbial taxa and functions in the mouse gut microbiome. (a–b) Peak-time distributions of functional genes (KOs) (a) and species (b) across three mice. Each dot represents the mean peak time of a feature, and lines connect the same features across individuals. Pairwise Jammalamadaka–SenGupta circular correlation coefficients (*ρ*_*c*_) between mice are indicated. Values approaching 1 indicate a strong positive circular association, whereas values near 0 indicate little circular association; all reported correlations were significant (*P* < 0.0001). Light and dark phases are indicated by white and grey backgrounds, respectively. (c) Peak-time density distributions of selected functional gene (KO) sets for each mouse. Curves represent kernel density estimates of peak times for KOs grouped by functional category. KO included each gene category shown in Additional file 2: Table S23. (d) Peak-time density distributions of selected microbial taxa for each mouse. Curves represent kernel density estimates of peak times for MAGs belonging to each taxonomic group. Dashed curves indicate the peak-time density distributions of all KOs assigned to the light phase (yellow) or dark phase (purple).

During the light phase, gene subsets exhibited a sequential peak-phase pattern. Genes involved in peptidoglycan biosynthesis and PTS-mediated carbon uptake (“Nutrient Acquisition” in Fig. 5c) peaked early, whereas those associated with rapid proliferation, including ribosome biogenesis, DNA replication, and aminoacyl-tRNA synthesis (“Rapid Proliferation”), as well as those related to respiratory metabolism, including electron transport chain components and the biosynthesis of key cofactors such as NAD, riboflavin, folate, and CoA (“Respiratory Metabolism”), reached maximal abundance later in the phase (Fig. 5c).

During the dark phase, a similar temporal progression was observed. Genes related to sulfur metabolism and glutathione biosynthesis (Sulfur & Redox Recovery) peaked early, followed by those involved in peptidoglycan-derived GlcNAc salvage and membrane lipid remodeling (GlcNAc Scavenger), and subsequently by genes associated with motility and active foraging, including flagellar assembly, chemotaxis, and broad-spectrum ABC transporters (Motile Active Forager), which reached their maxima later in the phase.

Taxonomic groups at the species level also exhibited distinct peak-time distributions across the circadian cycle (Fig. 5d). *Limosilactobacillus reuteri* and *Lactobacillus johnsonii* reached peak abundance during the early light phase, whereas *Muribaculaceae* and *Bacteroidaceae* peaked later within the light phase. In contrast, *Lachnospiraceae* and *Ligilactobacillus murinus* showed peak densities predominantly during the dark phase, with *Lachnospiraceae* reaching peak abundance earlier than *Ligilactobacillus murinus*.

The peak-phase ordering of these taxa and functional gene categories were preserved across mice, indicating that the within-phase sequence is a reproducible feature of microbiome temporal organization rather than an individual-specific pattern. Moreover, the peak-time ordering of both microbial taxa and functional gene categories remained stable throughout the observation period (see Additional file 1: Fig. S16b–c).

## Discussion

In this study, we show that the mouse gut microbiota exhibits reproducible phase-synchronized temporal organization at the community level. Although previous studies have reported compositional differences between day and night [1,4], our high-resolution phase-based analysis extends this view by resolving temporal phase relationships among microbial taxa and functional genes and by tracking community-level phase coherence over time and following perturbation. Here, synchronization refers to observed phase alignment rather than to a specific underlying coupling mechanism.

This analysis was enabled by continuous automated fecal sampling, together with GORIN, an analytical framework that integrates full-length 16S rRNA amplicon sequencing with long-read metagenomics of pooled samples. This framework reconstructs genome- and function-level microbiome trajectories cost-effectively, without requiring per-sample shotgun metagenomics. Full-length 16S rRNA sequences recovered in the long-read MAGs allow amplicon variants to be directly linked to reconstructed genomes, enabling simultaneous tracking of taxonomic and functional dynamics. The automated sampling system provided the dense temporal resolution needed to resolve community changes as continuous trajectories, while genome-level resolution was maintained through a limited number of long-read reference datasets. This approach provides a scalable strategy for generating genome-resolved microbiome time series.

A notable finding of this study is the greater temporal stability of functional gene content relative to taxonomic composition. Whereas MAG-level profiles showed both multi-day drift and diurnal cycling, KO-level profiles were dominated by the diurnal cycle with little evidence of long-term divergence, indicating that the functional repertoire of the community is more tightly buffered than its taxonomic composition. This functional stability appears to arise through at least two distinct mechanisms. For highly redundant functions encoded by many genomes, statistical averaging across contributors produces smooth temporal trajectories analogous to the portfolio effect described in ecology [54]. For functions encoded by fewer genomes, stability is maintained through compensatory dynamics, whereby the identities of contributing taxa shift over time while their aggregate functional abundance remains relatively stable. Together, these mechanisms suggest that functional stability can emerge at the community level despite substantial taxonomic turnover. We note that these mechanisms were not tested directly, and that the 14-day observation period does not establish whether this stability would persist under stronger or more prolonged taxonomic perturbation.

Phase analysis, which targets the diurnal component rather than the multi-day drift, separated the community into two groups that peak during the light and dark phases, respectively, and that differ in gene content. This functional differentiation characterizes, at the genome level, the metabolic potential associated with this temporal partitioning. While prior studies have documented shifts in the relative abundance of *Bacteroidota* and *Bacillota* across the light-dark cycle [1,4], the present study shows that these compositional shifts are accompanied by distinct carbohydrate-active gene repertories at the level of individual genomes. Because gene content is strongly associated with microbial lineage, we repeated the functional comparisons within individual phyla. This removed the enrichment of Gram-negative cell-envelope functions, consistent with these being lineage markers, but motility and several carbohydrate-related differences persisted. In particular, Dark-associated trehalose biosynthesis and starch- and glycogen-related CAZyme families were enriched within both *Bacillota_A* and *Bacillota*, indicating a phase-associated functional bias shared across two separate phyla. Several Light-associated glycan-degrading functions also remained significant within *Bacteroidota* and showed the same direction in another phylum, although none reached significance in more than one phylum.

The enrichment of mucosal glycan-degrading CAZyme families in the Light cluster is particularly noteworthy in the context of intestinal transit dynamics. Mice feed predominantly during the dark phase, and whole gut transit is substantially faster during the active phase (approximately 2.5 hours) than during the rest phase (approximately 4 hours) [56]. In line with this, our high-resolution fecal output data showed that pellet production was markedly higher during the dark phase than the light phase (Fig. 1e), consistent with greater intestinal throughput during the dark phase. The light-phase community, corresponding to the host resting phase when feeding activity is reduced, is thus enriched in mucosal glycan-degrading rather than dietary polysaccharide-degrading CAZyme families. These analyses describe the metabolic potential encoded in the reconstructed genomes rather than measured activity. Establishing whether these differences in genomic potential are realized as phase-dependent differences in metabolic activity will require transcriptomic or metabolomic measurements.

Our high-resolution time series further revealed ordered diel succession of microbial taxa and metabolic functions within each circadian phase (Fig. 5). Functional genes displayed conserved peak-time orderings across individual mice, indicating reproducible temporal organization of functional gene content across individuals. More broadly, such temporally organized functional succession may shape host physiology and ecosystem-level functions, raising the possibility that disruption of microbial rhythmicity contributes to dysbiosis.

The perturbation experiments provide direct evidence that phase coherence in the gut microbiota is a dynamical property subject to disruption and recovery. This analysis revealed a transient loss of coherence immediately following cage transfer, followed by a gradual and coordinated recovery as the host acclimated. Perturbation experiments further showed that antibiotic treatment also disrupted this coherence, after which phase coherence gradually recovered over the following days. Microbial dynamics in the gut are constrained by strong external drivers, including host circadian rhythms and feeding cycles. Whether the coherence measured here arises from interactions among taxa or from their independent entrainment to these shared drivers cannot be determined from the present data, since both would produce the same loss and recovery; the recovery timescale may additionally reflect declining antibiotic exposure and subsequent ecological regrowth. Applying this approach in experiments that perturb host circadian regulation, as well as in disease models, may help distinguish these mechanisms and clarify their physiological consequences.

A further observation bearing on the temporal organization of the community is that fecal peak times were consistently positioned between the Light and Dark cluster distributions, at the phase boundary separating the two community states (Fig. 4b). This pattern was preserved day-to-day across the 14-day observation period in both mice examined. Under the fixed 12:12 light–dark cycle used here, fecal excretion timing and Zeitgeber time co-vary, so the present design cannot determine whether community phase is more closely aligned with fecal excretion timing or with Zeitgeber time. This association therefore motivates the hypothesis that microbial community dynamics are coupled to host gut physiology, although the present data do not establish such coupling.

The present study has several limitations. First, our conclusions are based on observations of six mice, with continuous recordings of at least two weeks obtained from four individuals, and the antibiotic perturbation experiment was performed in a single animal. Although the key dynamical features were reproducible across animals, larger studies will be necessary to more comprehensively assess inter-individual variability and the generalizability of these findings, including potential sex-related differences: only female mice were examined here, and sex differences have been reported both in murine gastrointestinal physiology [57] and in the rhythmicity of the intestinal microbiota [3].

Second, our analysis relies on microbial abundances measured in fecal samples and expressed per milligram of feces. As such, the observed temporal structures do not directly reflect spatial heterogeneity along the gastrointestinal tract. In addition, because gastrointestinal transit and fecal output themselves exhibit circadian variation in mice [56], part of the observed diurnal amplitude may reflect changes in excretion rate and dilution rather than in microbial population size within the gut. The observation of the same bimodal structure in spike-in-normalized absolute abundances (Additional file 1: Fig. S12) argues against a purely compositional origin of this pattern, but does not exclude such effects. Consequently, the precise correspondence between the detected microbial phase dynamics and specific stages of host physiology remains constrained.

Third, genome- and function-level trajectories were inferred from amplicon counts rather than measured directly. Incomplete MAGs may lack functional genes or 16S rRNA gene copies, potentially leading to underestimation of functional repertoires or overestimation of MAG abundance, respectively. Shared ASVs and potential assembly or binning errors may also introduce uncertainty in abundance allocation among MAGs and consequently in reconstructed functional trajectories. Although comparison with short-read metagenomes supports the inferred functional composition overall, it does not exclude effects on individual functions. The absence of an annotation should therefore not be interpreted as biological absence.

Despite these limitations, the experimental platform developed here offers substantial opportunities for future expansion. As fecal samples are immediately preserved in RNA*later*™, ensuring high nucleic acid stability, the system is readily adaptable to analyses beyond those performed in this study, including phage dynamics and host transcriptomics. Because the reconstruction framework requires only a limited number of long-read references, it could also be extended to human cohorts, in which dense longitudinal studies have shown substantial fluctuations around relatively stable individual-specific community configurations [58–61]. In such cohorts, this framework could be used to track changes in community coherence across antibiotic exposure, dietary transitions, or circadian disruption. Disruption of microbial rhythmicity has in turn been associated with metabolic dysfunction [1,2]. Applied more broadly, the framework introduced here may help clarify the functional consequences of disrupted microbial temporal organization in the context of host health and disease.

## Conclusions

This study reveals fine-scale temporal organization in the gut microbiome. We identified two phase-coherent groups with distinct encoded gene repertoires and reproducible temporal ordering across individuals. Phase coherence was transiently disrupted following perturbation and recovered over several days, demonstrating that these relationships can be quantitatively tracked over time. The automated sampling platform, genome-resolved profiling approach, and phase-analytic tools introduced here provide a scalable foundation for investigating how microbial temporal organization shapes host physiology and contributes to dysbiosis.

## Declarations

## Ethics approval and consent to participate

All animal experiments were approved by the Animal Experiments Committee of RIKEN (approval number AEY2024-32).

## Consent for publication

Not applicable.

## Availability of data and material

The datasets supporting the conclusions of this article are available in the DNA Data Bank of Japan (DDBJ) repository, under BioProject accession numbers PRJDB37572 and PRJDB39835.

The custom scripts used for data processing and analysis are publicly available at the following repositories:

GORIN: https://github.com/musashi021203/GORIN

NPPD: https://github.com/rie-maskawa/Non-parametric-Period-Detection

## Authors’ contributions

R.K., R.M., W.S., L.T. and M.Takayasu conceived the study. R.K. and R.M. curated the data. Formal analyses were performed by R.K., R.M., M.A. and H.M. Funding was acquired by W.S., M.Takayasu and L.T. Investigations were carried out by R.K., C.S., M.Tanokura, M.Takagi and K.K. Methodology was developed by R.M., M.A., Y.Y., T.R., H.T., L.T., M.Takayasu, R.K., H.M. and W.S. Software was developed by R.M., M.A., H.T. and M.Takayasu. The study was supervised by W.S. and M.Takayasu. R.M. and R.K. prepared the visualizations. The original draft was written by R.K., R.M., H.T., M.Takayasu and W.S., and all authors, including I.D.V. and L.T., contributed to reviewing and editing the manuscript.

## Acknowledgments

We thank the Animal Facility staff at RIKEN Yokohama for accommodating the installation of our experimental device in the animal room and for providing routine animal husbandry and care. We also gratefully acknowledge Advanced U-corporation Co., Ltd., for their collaboration and technical support in the development of the automated fecal collection device. Computational resources were provided by the HOKUSAI BigWaterfall2 supercomputing system at RIKEN. We also acknowledge the Genome and Transcriptome Unit (GTU) at RIKEN for high-throughput sequencing. We also thank Editage (www.editage.jp) for English language editing. We appreciate helpful discussions and logistical support from colleagues and laboratory members throughout the course of this study.

## List of abbreviations

ASV: Amplicon sequence variant
CAZyme: Carbohydrate-active enzyme
MAG: Metagenome-assembled genome
HMQ-MAG: High- and medium-quality metagenome-assembled genome
KO: KEGG Ortholog
GORIN: Genome abundance Optimization from 16S rRNA-Informed Networks
NPPD: Non-parametric Period Detection
PCoA: Principal coordinates analysis
SPF: Specific-pathogen-free
ZT: Zeitgeber time

## Additional files

Additional file 1: Supplementary Information.

File format: .docx

Title of data: Supplementary Information.

Description of data: This file contains Supplementary Notes, Supplementary Figs. S1–S16, Supplementary Tables S3, and titles and descriptions for Supplementary Tables S1, S2, S4– S23.

Additional file 2: Supplementary Tables (Table S1, S2, S4–Table S23)

File format: .xlsx

Title of data: Supplementary tables supporting the study.

Description of data: This file contains Tables S1, S2, S4–Table S23, which include detailed datasets supporting the analyses presented in the manuscript. Individual table titles and descriptions are provided in Additional file 1.

