## Supplementary info for "High-resolution temporal profiling reveals synchronized dynamics of the mouse gut microbiome"

**Additional file1: Supplementary Information**

* Corresponding authors:

**Table of Contents**

Supplementary Notes

1. Device architecture and mechanical design of the automated mouse fecal collection system
2. Validation of the automated fecal collection workflow

Supplementary Tables

Table S1. Mouse characteristics and experimental design.

Table S2. Composition of the long-read sequencing pools.

Table S3. Identifiability of MAG abundance estimates and cross-taxonomic ASV sharing in each mouse.

Table S4. ASV–MAG correspondence matrix C for Mouse A.

Table S5. ASV–MAG correspondence matrix C for Mouse B.

Table S6. ASV–MAG correspondence matrix C for Mouse C.

Table S7. ASV–MAG correspondence matrix C for Mouse D.

Table S8. ASV–MAG correspondence matrix C for Mouse E.

Table S9. ASV–MAG correspondence matrix C for Mouse F.

Table S10. List of representative MAGs.

Table S11. MAG–Gene matrix G (KEGG Orthology annotations) for Mouse A.

Table S12. MAG–Gene matrix G (KEGG Orthology annotations) for Mouse B.

Table S13. MAG–Gene matrix G (KEGG Orthology annotations) for Mouse C.

Table S14. MAG–Gene matrix G (KEGG Orthology annotations) for Mouse D.

Table S15. MAG–Gene matrix G (KEGG Orthology annotations) for Mouse E.

Table S16. MAG–Gene matrix G (KEGG Orthology annotations) for Mouse F.

Table S17. MAG–Gene matrix G (CAZyme annotations) for Mouse A.

Table S18. MAG–Gene matrix G (CAZyme annotations) for Mouse B.

Table S19. MAG–Gene matrix G (CAZyme annotations) for Mouse C.

Table S20. MAG–Gene matrix G (CAZyme annotations) for Mouse D.

Table S21. MAG–Gene matrix G (CAZyme annotations) for Mouse E.

Table S22. MAG–Gene matrix G (CAZyme annotations) for Mouse F.

Table S23. KEGG orthologs (KOs) with peak-phase values across mice and associated functional annotations used in Fig. 5C.

Supplementary Figures

Fig. S1 Supplementary optimization analysis for MAG–ASV abundance reconstruction.

Fig. S2 Validation of inferred taxonomic and functional profiles using short-read shotgun metagenomics.

Fig. S3 Non-parametric Period Detection (NPPD) for identifying rhythmic dynamics in microbiome time series.

Fig. S4 Minute-scale temporal profiling of the mouse gut microbiome using an automated fecal collection system.

Fig. S5 Validation of the automated fecal collection system against manual collection.

Fig. S6 Bipartite graph between ASVs and MAGs based on genome correspondence.

Fig. S7 High-resolution longitudinal profiles of gut microbiome composition across multiple mice.

Fig. S8 Multiscale temporal dynamics of taxonomic and functional composition.

Fig. S9 Examples of complex temporal dynamics observed in the metagenomic time-series data.

Fig. S10 NPPD resolves time-varying local phase that is not represented by a single global period and phase.

Fig. S11 Core oscillatory MAGs and phase difference structure.

Fig. S12 Temporal dynamics and phase behavior of absolute MAG abundances.

Fig. S13 Taxonomic and functional characteristics of Light and Dark clusters across mice.

Fig. S14 Functional comparison between the Light and Dark clusters within individual phyla.

Fig. S15 Phase-based results are robust to the valid-cycle criterion used to define core oscillatory MAGs.

Fig. S16 Phase organization of functional and taxonomic dynamics across mice.

**Supplementary Notes**

***1. Device architecture and mechanical design of the automated mouse fecal collection system***

*Overall configuration*

The automated fecal collection device consists of three integrated modules:

(1) a metabolic cage that separates feces and urine by gravity and geometry,

(2) a fraction collection module that houses individual sampling tubes, and

(3) a control and actuation unit that regulates tube indexing and sealing.

These modules are arranged in a vertical configuration to allow unobstructed free fall of pellets while keeping all mechanical components below the animal-housing compartment.

*Metabolic cage and separation funnel*

The metabolic cage (TECNIPLAST S.p.A., Italy) is equipped with a stainless-steel wire-mesh floor that allows feces and urine to pass through without retention. Beneath the floor, a conical separation funnel directs fecal pellets straight downward through the central opening, while urine flows along the funnel wall and is diverted toward a separate outlet via a dedicated guiding groove. No suction or active flow is applied, preserving natural excretion patterns.

*Fraction collection module*

The fraction collector consists of a circular tube rack in which 24 fecal tubes are arranged along the outer circumference and 24 urine tubes are positioned concentrically on the inner ring. Both fecal and urine tubes were 4.5-mL CryoTubes (Nalge Nunc, 92 × φ12.5 mm; Thermo Fisher Scientific). Each fecal tube was pre-filled with 3 mL of RNA*later*™ solution to stabilize the microbiome immediately upon pellet entry. The module is mounted directly below the funnel outlet to ensure that pellets enter the designated tubes solely by gravity. Tube indexing is controlled by a stepper-motor–driven rotation stage that advances the rack at fixed intervals (normally every 60 minutes, adjustable from 1 minute to several hours).

*Tube-sealing mechanism (stainless-steel ball system)*

The tube is sealed at each indexing interval by a stainless-steel ball delivered via a swing-arm mechanism positioned beneath the ball reservoir. The reservoir stores the balls in a spiral track that feeds them one by one into a central receiving hole on the swing arm. During operation, the swing arm rotates to align this receiving hole with the inlet of a plastic guide tube leading to either the fecal or urine collection tube, releases a single ball into the guide, and then returns to its central standby position where no ball is released. The ball travels through the guide tube and enters the corresponding collection tube, reducing evaporation and preventing cross-contamination between sequential samples.

*Control unit and timing program*

A control unit manages the rotation of the turntable and the operation of the ball-delivery mechanism.

The unit consists of a touch-panel interface and dedicated controllers for the indexing stage and the sealing mechanism. Through the touch panel, the user specifies the number of collection tubes and the switching interval, then initiates operation by pressing the start button. The control system includes a real-time clock for interval scheduling and timestamp display, as well as circuitry for turntable rotation and ball-delivery actuation. The control interval (sampling frequency) is user-programmable from 1 second to 99 hours. In the present study, hourly sampling was used for the time-series experiments. Sampling at shorter intervals was evaluated on day 4 in Mouse P1 and Mouse P2, with 24 collection intervals of 10 and 5 minutes, respectively, following 3 days of hourly sampling (Fig. S4).

*Operating stability and maintenance*

During extended operation, the device maintained stable mechanical performance, and no indexing misalignment occurred after initial calibration. Although the system operated automatically during each sampling interval, daily tube replacement and system restart were performed manually. A small number of fecal pellets were occasionally retained on the funnel surface or dropped outside the guide path and were therefore not collected. The metabolic-cage module can be removed easily. The cage portion is autoclavable and was cleaned according to the animal facility's standard protocol after each experimental run. During operation, the mouse-housing compartment was wiped but not washed to avoid disturbing the animals, and the separation funnel was rinsed with water as needed.

***2. Validation of the automated fecal collection workflow***

*Validation design*

We evaluated whether the automated collection workflow, including sample preservation under routine operating conditions, yielded microbiome profiles comparable to those obtained by manual collection followed by immediate DNA extraction. Eight matched pairs were collected from Mouse V1–V4 at two sampling times, 14:00 (ZT7) and 19:00 (ZT12), as described in the Methods. Within- and between-mouse variation provided a biological context for interpreting the differences between collection workflows.

*Comparison of microbial profiles*

Relative abundances were compared at the ASV, species, and genus levels (Fig. S5a). Species- and genus-level profiles were obtained by aggregating ASV abundances according to their reference assignments at nucleotide identity thresholds of ≥97% and ≥94%, respectively. Spearman's correlations were calculated separately for each matched pair at each resolution, including features with zero abundance in one or both samples. The median coefficient across the eight pairs was reported. To contextualize differences between collection workflows, samples from Mouse E, Mouse F, Mouse P1, and Mouse P2 were used to assess within- and between-mouse variation (Fig. S5b). The selected 24-h period immediately followed the first 48 h of collection and contained 21, 19, 21, and 21 samples, respectively. Within-mouse comparisons included all pairs of samples collected at different time points from the same mouse (801 pairs). Between-mouse comparisons included all combinations of samples from different mice within this period, without restriction to the same time of day (2,520 pairs).

For ASV-level Bray–Curtis distance calculations, each library was subsampled without replacement to 10,000 reads in each of 100 iterations. Counts were converted to relative abundances, and pairwise Bray–Curtis distances were calculated separately in each iteration and averaged across iterations. Reproducible random seeds derived from library identifiers ensured consistent subsampling of the same libraries across comparisons. Distances were summarized descriptively for the eight matched automated–manual pairs and the within- and between-mouse comparisons. No inferential statistical tests were applied to these pairwise distances.

The 16 automated and manually collected validation samples were subjected to average-linkage hierarchical clustering using the mean ASV-level Bray–Curtis distances calculated above (Fig. S5c). Genus-level compositions were displayed, with the 12 assigned genera having the highest mean relative abundance across the 16 samples shown individually. Remaining assigned genera were pooled as Other, and sequences without a genus assignment were shown separately as Genus undefined. These aggregations were used only for visualization and did not affect the ASV-level distances used for clustering.

*Validation results*

The median Spearman correlation across the eight matched pairs was 0.724 at the ASV level, 0.906 at the species level, and 0.946 at the genus level (Fig. S5a). The median ASV-level Bray–Curtis distance between collection workflows was lower than those for the within- and between-mouse comparisons (Fig. S5b). In hierarchical clustering, each automatically collected sample clustered first with its matched manually collected sample in all eight pairs, and genus-level compositions were similar within pairs (Fig. S5c). These results support the ability of the automated workflow to capture fecal microbiome composition comparable to that obtained by manual collection under the conditions examined.

**Supplementary Tables**

*Supplementary Tables S1,S2 and S4–S23 are provided as separate Excel files.*

**Table S1. Mouse characteristics and experimental design.** Female C57BL/6J mice were used: Mouse A–F for the time-series experiments, and Mouse P1, Mouse P2, and Mouse V1–V4 for validation of the collection system. Samples from Mouse E and Mouse F were also used for collection-system validation. The table summarizes strain, sex, age, treatment conditions, sampling start time, administration timing, and sampling duration. Mouse C and Mouse D received a single oral gavage on day 4 (sterile water for C and vancomycin for D).

**Table S2. Composition of the long-read sequencing pools.** For Mouse A–D, each of the four pools per mouse combines the hourly samples of one phase from one part of the collection period. For Mouse E and Mouse F, samples from all time points over the four-day collection period were combined into a single pool per mouse. Pool indicates the sample ID assigned to each pooled sample. Day_first and Day_last indicate the first and last sampling days represented in each pool, respectively, with days numbered from the start of collection.

**Table S3. Identifiability of MAG abundance estimates and cross-taxonomic ASV sharing in each mouse.** The ASV–MAG bipartite network was partitioned into connected components, each defining a local system of linear equations. MAGs and ASVs are termed identifiable when they belong to a component whose copy-number submatrix is of full column rank, so that the corresponding abundances are uniquely determined. Cross-genus and cross-family entries include only quality-suspect or unresolved MAGs that directly shared an ASV with a MAG assigned to a different genus or family, respectively, within rank-deficient components. Each MAG was counted once per category; cross-family targets are a subset of cross-genus targets. Abundance values are percentages of total inferred MAG abundance, summarized as medians (interquartile ranges) across time points. These values describe the abundance of the selected MAGs, not misallocation rates. Mouse E and Mouse F had no rank-deficient components and therefore no eligible targets for the cross-taxonomic assessment.

| | Mouse A | Mouse B | Mouse C | Mouse D | Mouse E | Mouse F |
|---|---|---|---|---|---|---|
| Components, n | 156 | 181 | 148 | 192 | 117 | 90 |
| Rank-deficient components, n | 19 | 30 | 24 | 28 | 0 | 0 |
| Identifiable MAGs / total | 221 / 329 | 274 / 463 | 212 / 340 | 302 / 456 | 157 / 157 | 119 / 119 |
| Identifiable ASVs / total | 315 / 401 | 365 / 527 | 296 / 391 | 419 / 554 | 288 / 288 | 241 / 241 |
| Identifiable MAG abundance, % | 72.7 (66.5–76.9) | 68.8 (62.9–76.2) | 65.4 (60.8–70.2) | 67.6 (63.1–72.6) | 100.0 (100.0–100.0) | 100.0 (100.0–100.0) |
| Cross-genus MAGs, n | 9 | 16 | 4 | 16 | 0 | 0 |
| Cross-genus abundance, % | 3.08 (1.85–4.15) | 1.33 (0.85–1.95) | 0.84 (0.56–1.22) | 0.82 (0.42–1.32) | 0 (0–0) | 0 (0–0) |
| Cross-family MAGs, n | 1 | 5 | 3 | 2 | 0 | 0 |
| Cross-family abundance, % | 0.026 (0.000–0.199) | 0.078 (0.033–0.162) | 0.420 (0.232–0.683) | 0.132 (0.060–0.232) | 0 (0–0) | 0 (0–0) |

**Table S4. ASV–MAG correspondence matrix C for Mouse A.** This dataset contains the correspondence matrix C linking amplicon sequence variants (ASVs) to High- and Medium- quality MAGs (HMQMAGs) reconstructed from Mouse A. The table was used to integrate ASV- and MAG-level analyses throughout the study.

**Table S5. ASV–MAG correspondence matrix C for Mouse B.** This dataset contains the correspondence matrix C linking ASVs to HMQMAGs reconstructed from Mouse B. The table was used to integrate ASV- and MAG-level analyses throughout the study.

**Table S6. ASV–MAG correspondence matrix C for Mouse C.** This dataset contains the correspondence matrix C linking ASVs to HMQMAGs reconstructed from Mouse C. The table was used to integrate ASV- and MAG-level analyses throughout the study.

**Table S7. ASV–MAG correspondence matrix C for Mouse D.** This dataset contains the correspondence matrix C linking ASVs to HMQMAGs reconstructed from Mouse D. The table was used to integrate ASV- and MAG-level analyses throughout the study.

**Table S8. ASV–MAG correspondence matrix C for Mouse E.** This dataset contains the correspondence matrix C linking ASVs to HMQMAGs reconstructed from Mouse E. The table was used to integrate ASV- and MAG-level analyses throughout the study.

**Table S9. ASV–MAG correspondence matrix C for Mouse F.** This dataset contains the correspondence matrix C linking ASVs to HMQMAGs reconstructed from Mouse F. The table was used to integrate ASV- and MAG-level analyses throughout the study.

**Table S10. List of representative MAGs.** This dataset lists representative MAGs defined separately for each mouse (Mouse A–F), including assembler information, genome quality metrics, genome lengths, GTDB-Tk taxonomy, and the non-representative MAGs grouped under each representative MAG.

**Table S11. MAG–Gene matrix G (KEGG Orthology annotations) for Mouse A.** This dataset contains the MAG–Gene correspondence matrix G linking each representative MAG to its KEGG Orthology (KO)–annotated genes reconstructed from Mouse A. These tables were used for functional profiling and pathway-level analyses.

**Table S12. MAG–Gene matrix G (KEGG Orthology annotations) for Mouse B.** This dataset contains the MAG–Gene correspondence matrix G linking each representative MAG to its KEGG

Orthology (KO)–annotated genes reconstructed from Mouse B. These tables were used for functional profiling and pathway-level analyses.

**Table S13. MAG–Gene matrix G (KEGG Orthology annotations) for Mouse C.** This dataset contains the MAG–Gene correspondence matrix G linking each representative MAG to its KEGG Orthology (KO)–annotated genes reconstructed from Mouse C. These tables were used for functional profiling and pathway-level analyses.

**Table S14. MAG–Gene matrix G (KEGG Orthology annotations) for Mouse D.** This dataset contains the MAG–Gene correspondence matrix G linking each representative MAG to its KEGG Orthology (KO)–annotated genes reconstructed from Mouse D. These tables were used for functional profiling and pathway-level analyses.

**Table S15. MAG–Gene matrix G (KEGG Orthology annotations) for Mouse E.** This dataset contains the MAG–Gene correspondence matrix G linking each representative MAG to its KEGG Orthology (KO)–annotated genes reconstructed from Mouse E. These tables were used for functional profiling and pathway-level analyses.

**Table S16. MAG–Gene matrix G (KEGG Orthology annotations) for Mouse F.** This dataset contains the MAG–Gene correspondence matrix G linking each representative MAG to its KEGG Orthology (KO)–annotated genes reconstructed from Mouse F. These tables were used for functional profiling and pathway-level analyses.

**Table S17. MAG–Gene matrix G (CAZyme annotations) for Mouse A.** This dataset contains the MAG–Gene correspondence matrix G linking each representative MAG to its CAZyme-annotated carbohydrate-active enzyme genes reconstructed from Mouse A. These tables were used to assess glycan-degradation and carbohydrate-utilization potentials.

**Table S18. MAG–Gene matrix G (CAZyme annotations) for Mouse B.** This dataset contains the MAG–Gene correspondence matrix G linking each representative MAG to its CAZyme-annotated carbohydrate-active enzyme genes reconstructed from Mouse B. These tables were used to assess glycan-degradation and carbohydrate-utilization potentials.

**Table S19. MAG–Gene matrix G (CAZyme annotations) for Mouse C.** This dataset contains the MAG–Gene correspondence matrix G linking each representative MAG to its CAZyme-annotated carbohydrate-active enzyme genes reconstructed from Mouse C. These tables were used to assess glycan-degradation and carbohydrate-utilization potentials.

**Table S20. MAG–Gene matrix G (CAZyme annotations) for Mouse D.** This dataset contains the MAG–Gene correspondence matrix G linking each representative MAG to its CAZyme-annotated carbohydrate-active enzyme genes reconstructed from Mouse D. These tables were used to assess glycan-degradation and carbohydrate-utilization potentials.

**Table S21. MAG–Gene matrix G (CAZyme annotations) for Mouse E.** This dataset contains the MAG–Gene correspondence matrix G linking each representative MAG to its CAZyme-annotated carbohydrate-active enzyme genes reconstructed from Mouse E. These tables were used to assess glycan-degradation and carbohydrate-utilization potentials.

**Table S22. MAG–Gene matrix G (CAZyme annotations) for Mouse F.** This dataset contains the MAG–Gene correspondence matrix G linking each representative MAG to its CAZyme-annotated carbohydrate-active enzyme genes reconstructed from Mouse F. These tables were used to assess glycan-degradation and carbohydrate-utilization potentials.

**Table S23. KEGG orthologs (KOs) with peak-phase values across mice and associated functional annotations used in Fig. 5C.** The table includes KO identifiers, gene names, peak phase (ZT) for each mouse, mean peak phase and standard deviation across mice, and functional annotations (category, pathway, module, and description).

### Supplementary figures

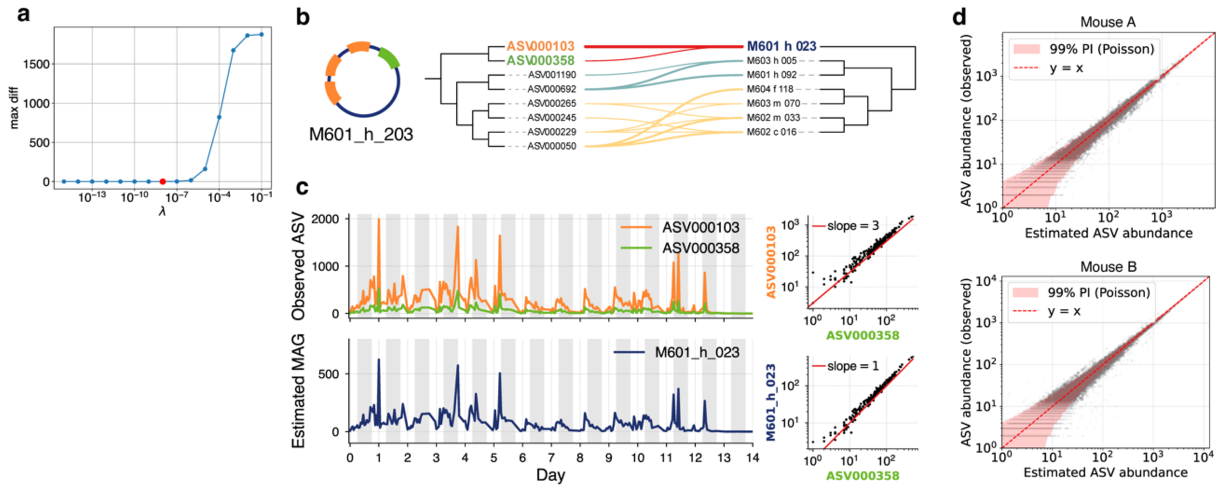


**Fig. S1** Supplementary optimization analysis for MAG–ASV abundance reconstruction. (a) Maximum difference in estimated MAG abundance, $\max_{i,t}|x_\lambda(i,t) - x_0(i,t)|$, as a function of the regularization parameter λ. Here, $x_0(i,t)$ denotes the reference abundance of MAG $i$ at time $t$ obtained under the baseline condition ($\lambda = 1.0 \times 10^{-8}$, $\varepsilon = 1.0 \times 10^{-4}$, $\delta = 1.0 \times 10^{-4}$), and $x_\lambda(i,t)$ denotes the abundance estimated with a different λ. (b) Example of a MAG (M601_h_023) containing multiple ASVs with different copy numbers. The circular diagram on the left represents the genome structure of the MAG, colored according to the associated ASVs. The right panel shows the correspondence between ASVs (left) and MAGs (right), arranged in the order of their phylogenetic trees. Edges belonging to the same connected component in the bipartite ASV–MAG network are shown in the same color. (c) Left panels show time series of two ASVs (top) and the corresponding estimated MAG abundance (bottom). Right panels show scatter plots of ASV abundances (top) and the relationship between the estimated MAG abundance and its constituent ASVs (bottom). (d) Relationship between observed and reconstructed ASV read counts for Mouse A and Mouse B. The horizontal axis represents the ASV read counts recalculated from the estimated MAG abundances, and the vertical axis represents the observed ASV read counts. Red shaded areas indicate the 99% Poisson prediction intervals computed based on observed counts.

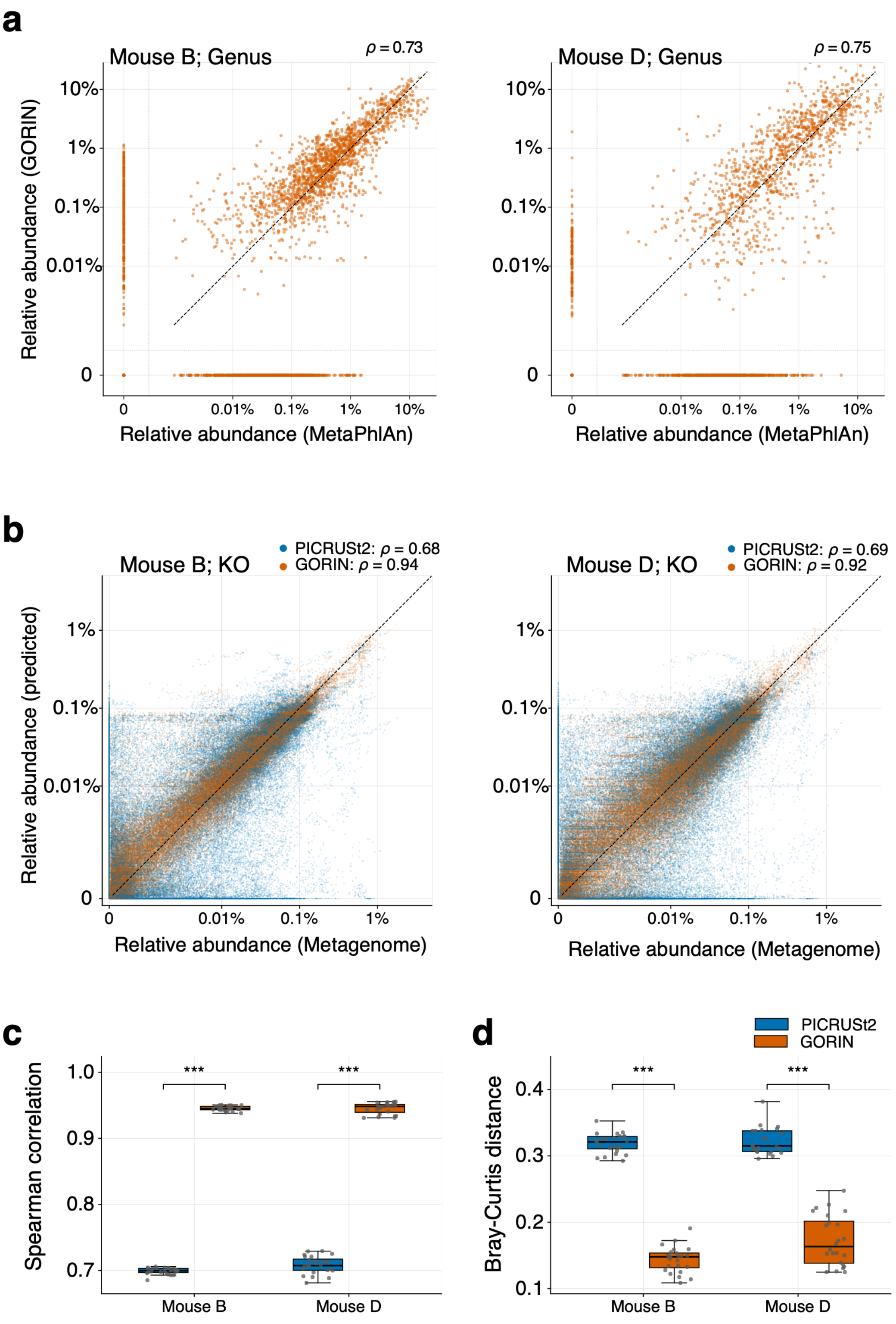


**Fig. S2** Validation of inferred taxonomic and functional profiles using short-read shotgun metagenomics. (a) Comparison of genus-level relative abundances between MetaPhlAn and GORIN for Mouse B (left) and Mouse D (right). The horizontal axis represents MetaPhlAn estimates, and the vertical axis represents inferred MAG abundances by GORIN aggregated at the genus level. Each point represents a genus at an individual sampling time. Spearman correlations are shown for all genus–

sample observations pooled across the 24 samples within each mouse. (b) Comparison of KO relative abundances between shotgun-derived profiles quantified using Salmon and profiles predicted using PICRUSt2 (blue) or GORIN (orange). Panels correspond to Mouse B (left) and Mouse D (right). Spearman correlations were calculated across all KO–sample observations within each mouse. In (a–b), dashed diagonal lines indicate equal relative abundances, and observations detected by only one method are shown on the corresponding zero axis. (c) Sample-wise Spearman correlations between predicted and shotgun-derived KO profiles. Colors indicate PICRUSt2 (blue) and GORIN (orange). Each point represents one sample (24 samples per mouse). Boxes indicate medians and interquartile ranges, and whiskers extend to the most extreme values within 1.5 times the interquartile range. (d) Sample-wise Bray–Curtis distances between predicted and shotgun-derived KO profiles, shown in the same format as (c). Prediction methods were compared using paired Wilcoxon signed-rank tests within each mouse ($***P < 0.001$).

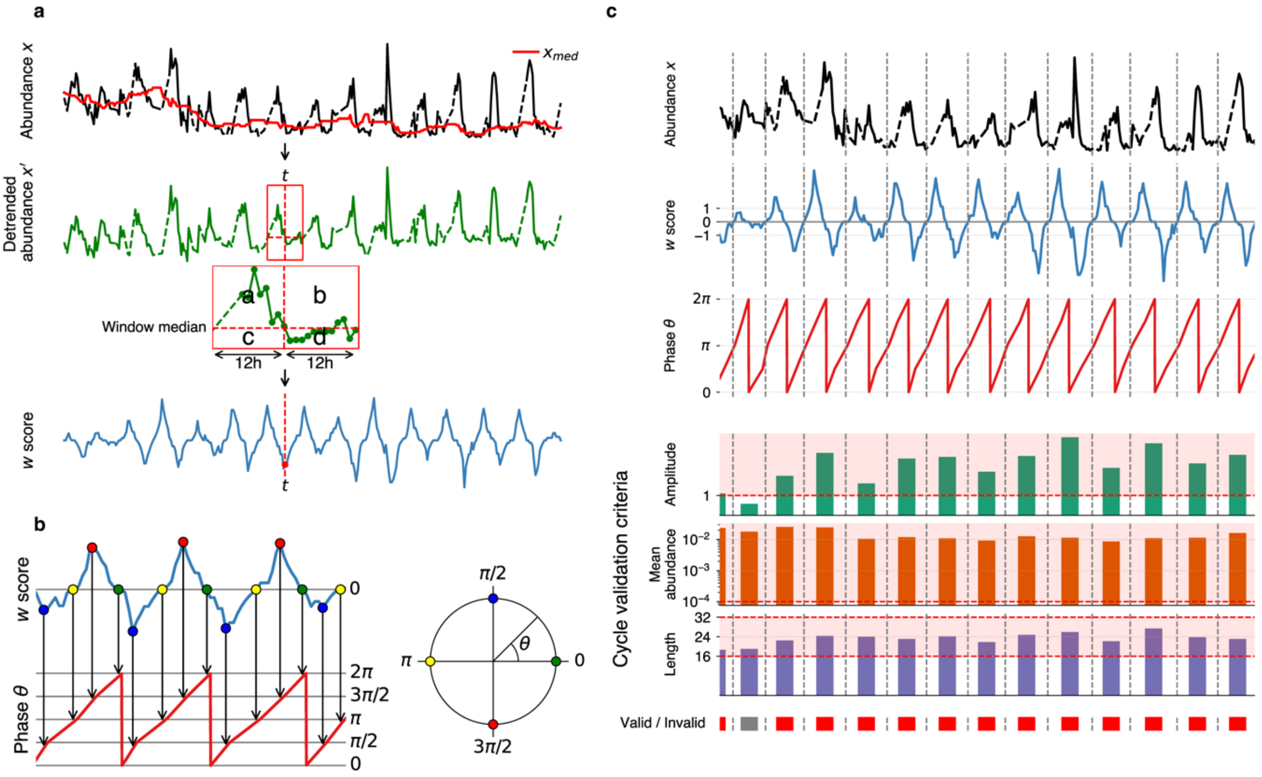


**Fig. S3** Non-parametric Period Detection (NPPD) for identifying rhythmic dynamics in microbiome time series. (a) Computation of the local significance score w(t). Top: the original abundance time series $x(t)$ (black) and its 24-hour moving median $x_{\mathrm{med}}(t)$ (red). Middle: the detrended signal. Bottom: the local significance score $w(t)$ calculated at each time point $t$ based on deviations in preceding and following 12-hour windows. (b) Phase assignment based on zero-crossings and local extrema of w(t). Oscillatory cycles are defined by successive zero-crossings and extrema, and used to construct a continuous phase trajectory, which is wrapped into the interval $[0,2\pi)$ to obtain the instantaneous phase $\theta(t)$. (c) Validation of detected oscillatory cycles. From top to bottom: original abundance, $w(t)$, phase, amplitude, duration, and cycle validation. Valid (red) and invalid (gray) cycles are shown at the bottom.

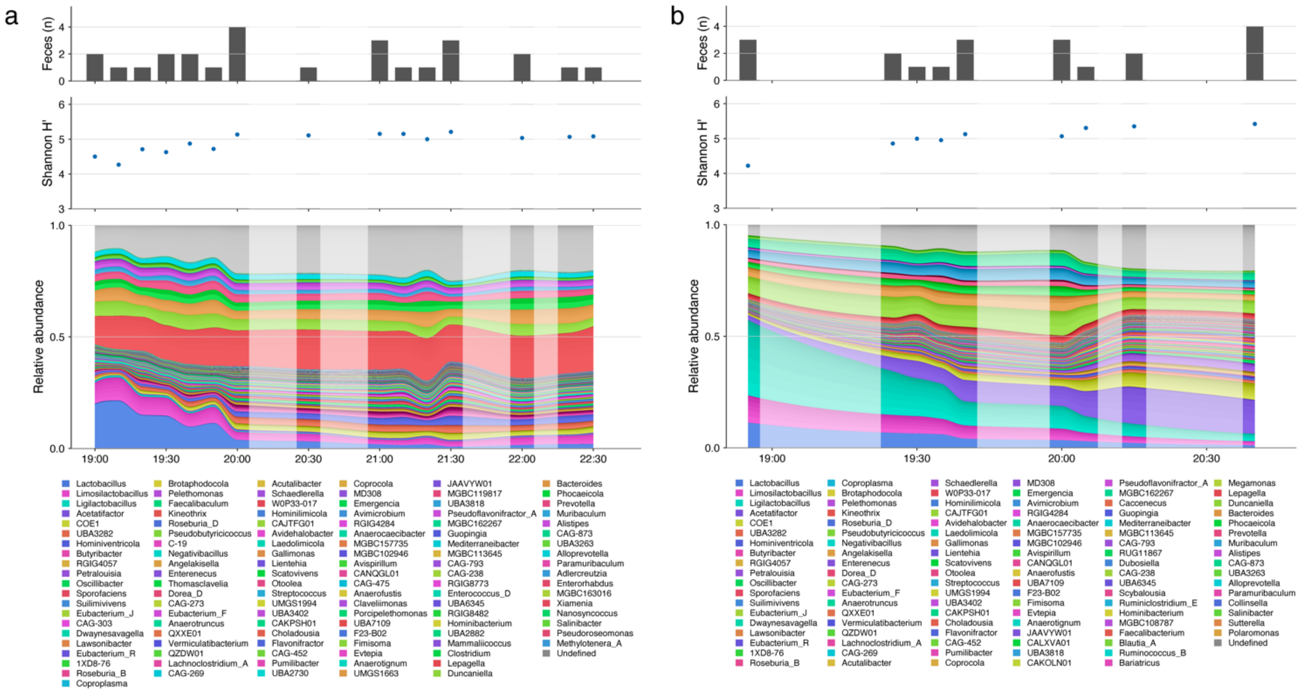


**Fig. S4** Minute-scale temporal profiling of the mouse gut microbiome using an automated fecal collection system. (**a**) Time-series microbiome data collected at 10-min sampling intervals. The upper panel shows the number of fecal pellets collected at each time point (gray bars). The middle panel shows Shannon diversity (H′) calculated for each sample (blue points). The lower panel shows the relative abundance of individual amplicon sequence variants (ASVs) as a stacked area plot. ASVs are grouped by phylum and genus. Colors indicate the genus of each ASV. Relative abundances were smoothed using piecewise cubic Hermite interpolating polynomial (PCHIP) interpolation. White vertical shading indicates time intervals for which no fecal sample was obtained. (**b**) Equivalent measurements obtained at 5-min sampling intervals. Panels are arranged as in (a).

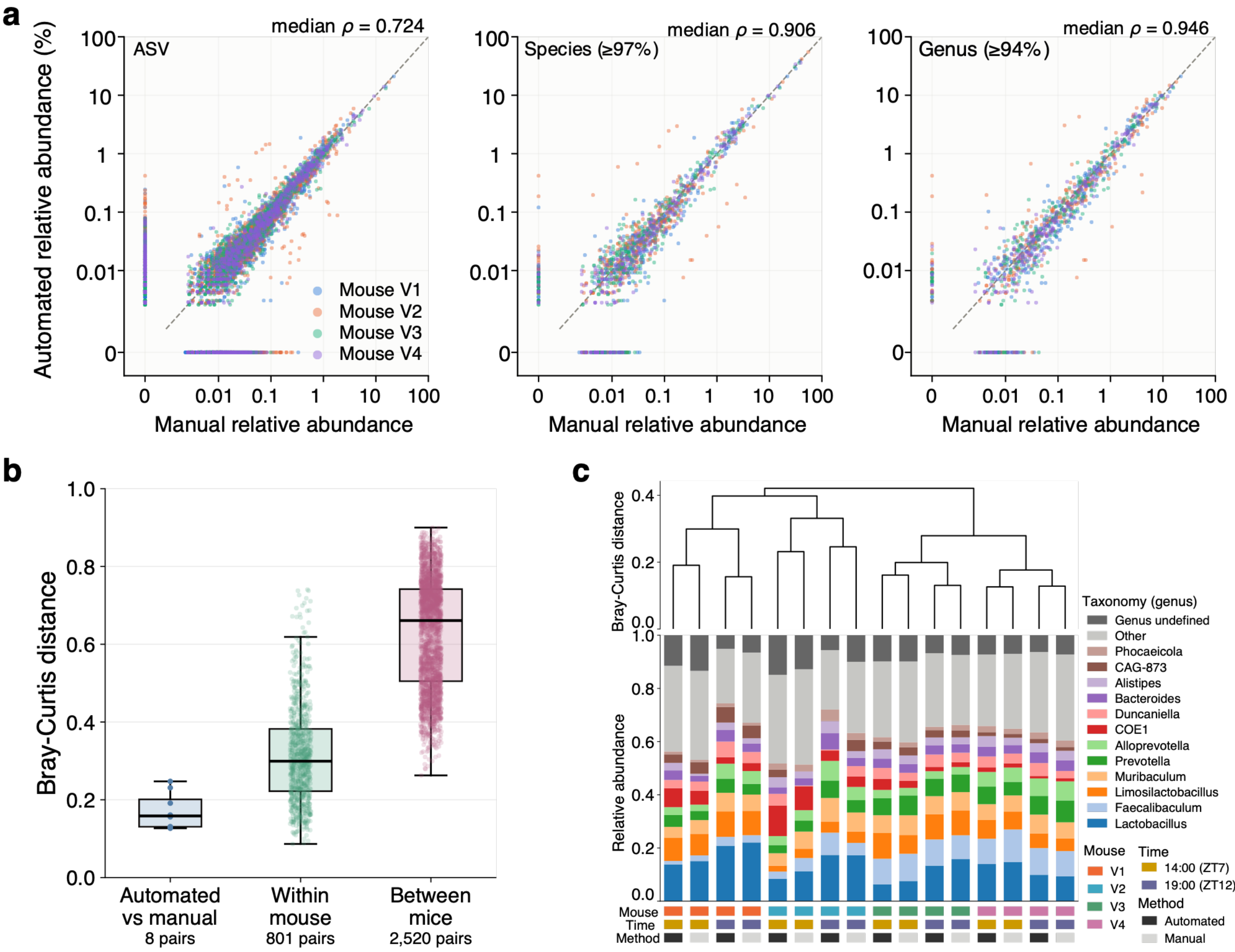


**Fig. S5** Validation of the automated fecal collection system against manual collection.

(a) Comparison of relative abundances between automated and manually collected pellets from eight matched pairs at the ASV, species, and genus levels. Each point represents one ASV, species, or genus in one matched pair. Axes are logarithmic; ASVs or taxa detected in only one member of a pair are plotted on the corresponding zero axis. Colors indicate mice; dashed lines indicate identity. Spearman's correlations were calculated separately for each matched pair; the median coefficient across the eight pairs is shown. (b) ASV-level Bray–Curtis distances between matched automated and manually collected pellets (8 pairs), between samples collected at different time points within the same mouse (801 pairs), and between samples from different mice (2,520 pairs). Within- and between-mouse comparisons used samples from Mouse E, Mouse F, Mouse P1, and Mouse P2 during a selected 24-h period. Points represent pairwise distances. Boxes indicate medians and interquartile ranges; whiskers extend to the most extreme values within 1.5 times the interquartile range. (c) Genus-level composition of the 16 validation samples, ordered by average-linkage hierarchical clustering based on ASV-level Bray–Curtis distances. The 12 genera with the highest mean relative abundance across samples are shown individually; remaining assigned genera are pooled as Other, and unassigned sequences are shown as Genus undefined. Annotation strips indicate mouse, sampling time, and collection method.

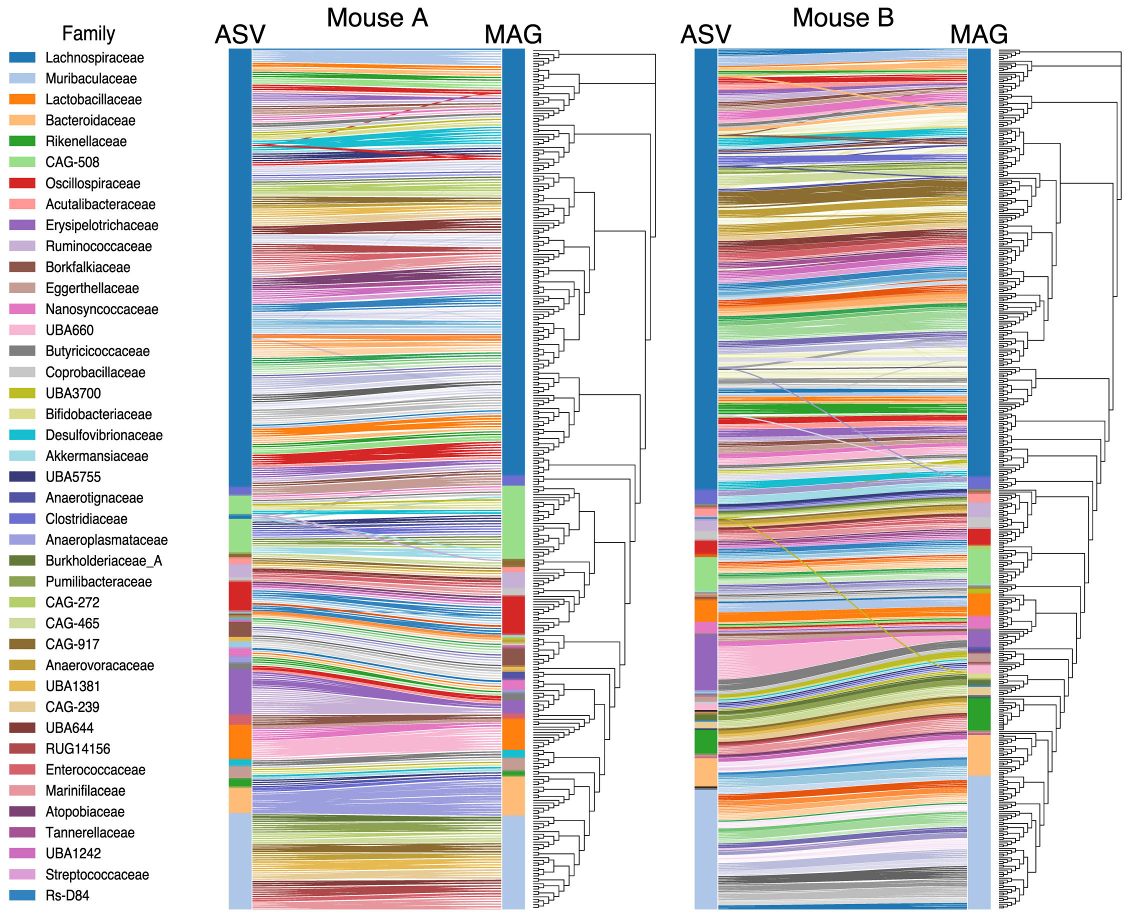


**Fig. S6** Bipartite graph between ASVs and MAGs based on genome correspondence. Each panel shows the mapping between ASVs and MAGs for Mouse A (left panel) and Mouse B (right panel). MAGs are ordered according to their phylogenetic relationships, while ASVs are sorted based on their correspondence to MAGs. Rows on each side are color-coded by family-level taxonomy independently assigned from the respective taxonomic reference databases used for ASVs and MAGs. Edges connect ASVs and MAGs belonging to the same connected component, with identical colors indicating one component. Most connected components were consistently annotated at the same family level, demonstrating the taxonomic coherence between ASV- and MAG-level reconstructions.

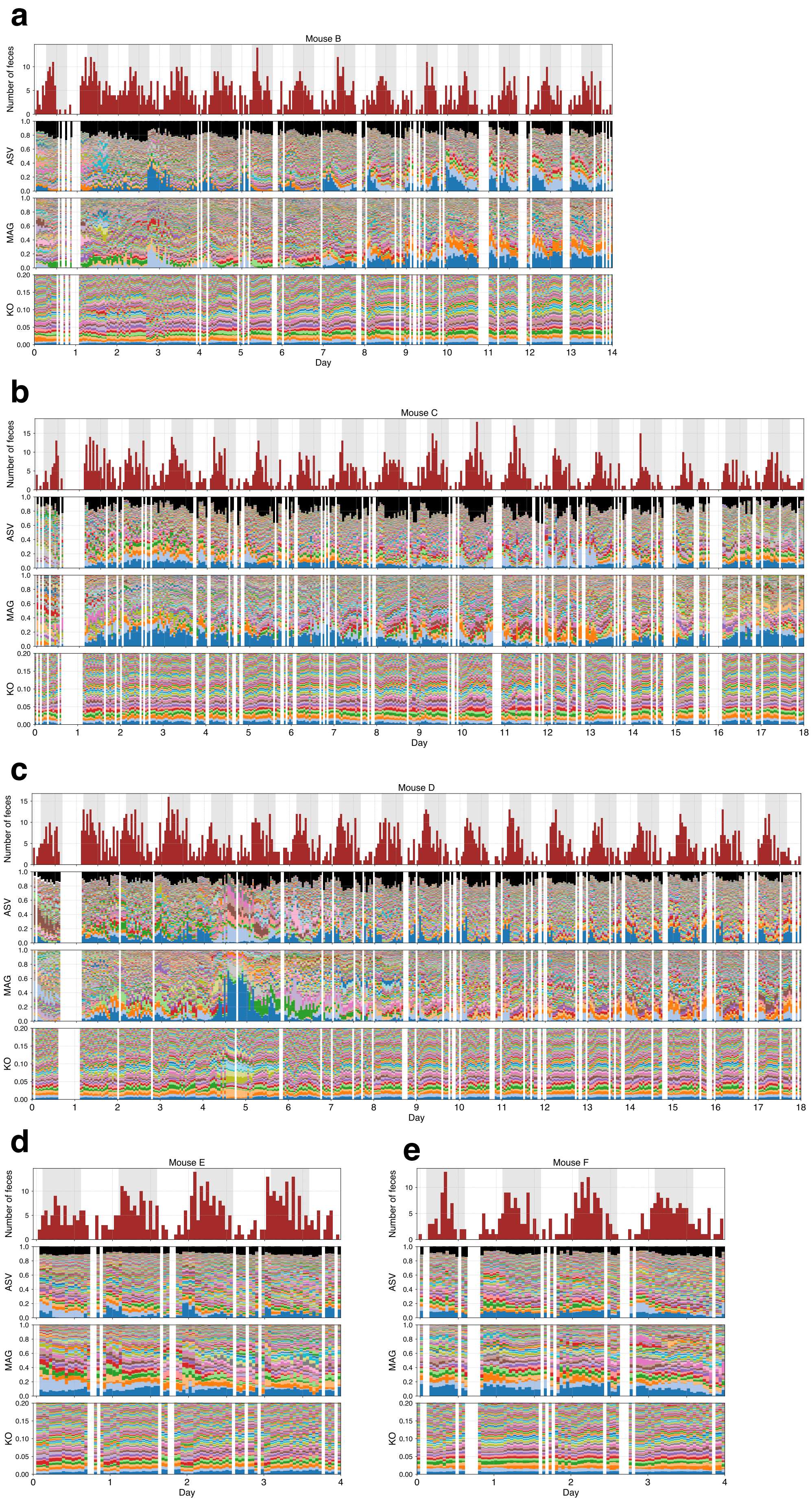
a
Mouse B
Number of feces
ASV
MAG
KO
Day
b
Mouse C
Number of feces
ASV
MAG
KO
Day
c
Mouse D
Number of feces
ASV
MAG
KO
Day
d
Mouse E
Number of feces
ASV
MAG
KO
Day
e
Mouse F
Number of feces
ASV
MAG
KO
Day

**Fig. S7** High-resolution longitudinal profiles of gut microbiome composition across multiple mice. From top to bottom in each panel: the number of fecal samples collected per time point, and the relative abundances of ASVs, MAGs, and KOs over time. Panels (a–e) correspond to individual mice (Mice B–F), each shown over its respective observation period. In the stacked plots, components are ordered from bottom to top according to their mean relative abundance. The KO panels are displayed with an expanded vertical scale (up to a relative abundance of 0.2) for visualization. Black regions in the ASV panels indicate ASVs not assigned to any MAG.

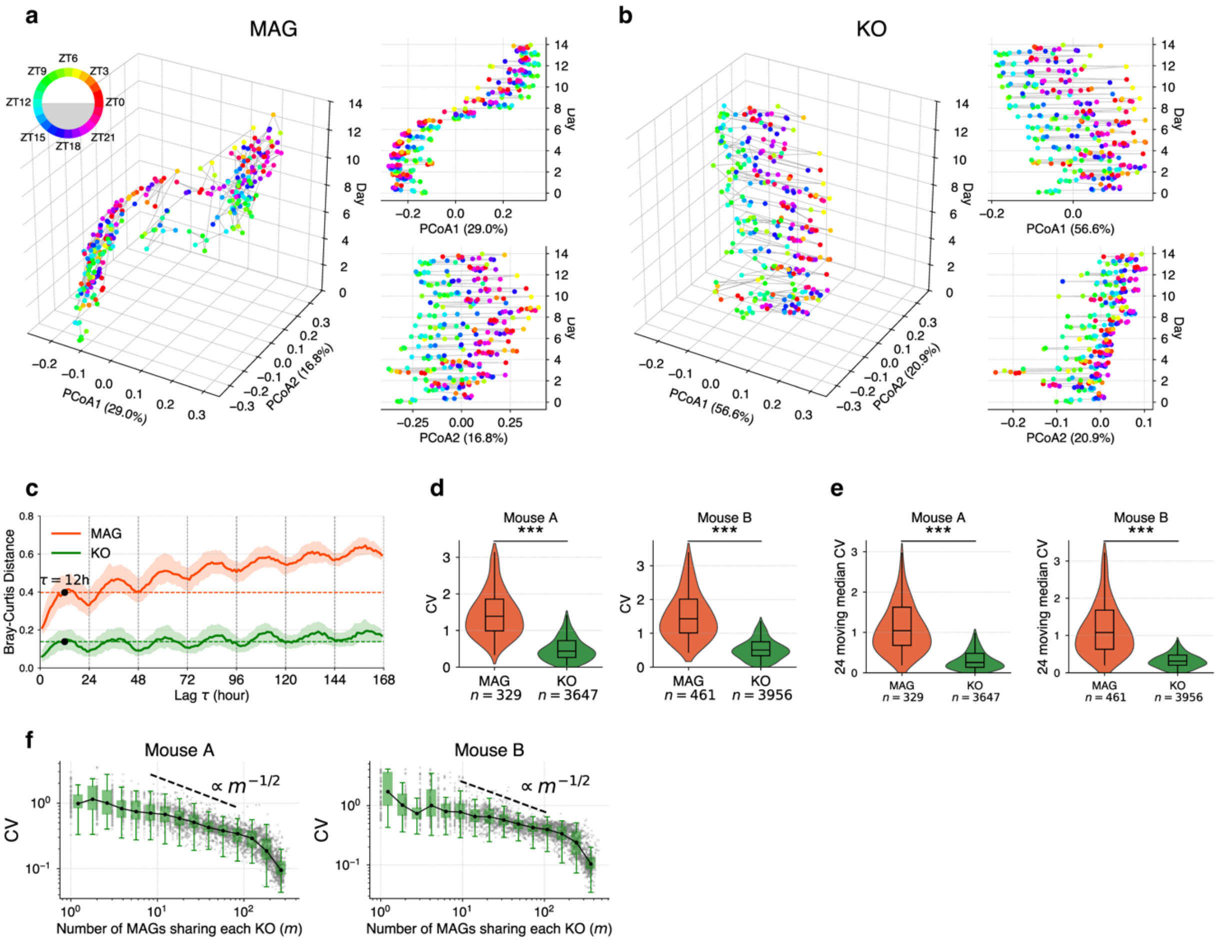


**Fig. S8** Multiscale temporal dynamics of taxonomic and functional composition. (a) PCoA of MAG-level compositions in Mouse B, based on Bray–Curtis distances. Each point represents a sample colored by sampling time (ZT). Left: three-dimensional plot with PCoA1–PCoA2 on the horizontal axes and time (day) on the vertical axis. Right: temporal trajectory of the principal coordinates. (b) PCoA of KO-level compositions in Mouse B, shown in the same format as (A). (c) Temporal Bray–Curtis distance as a function of time lag τ in Mouse B, illustrating the temporal autocorrelation structure of the community at the MAG (orange) and KO (green) levels. Solid lines indicate the median across all time windows, and shaded regions denote the interquartile range (IQR). (d) Violin plot of the coefficient of variation (CV) for MAG- and KO-level time series in Mice A and B, showing that functional (KO-level) profiles fluctuate less than taxonomic (MAG-level) profiles. ***$P < 0.001$, two-sided Mann–Whitney U test. (e) Violin plot showing the CV of 24-hour moving medians of relative abundance for MAG- and KO-level time series in Mice A and B. (f) Relationship between the number of MAGs sharing each KO (m) and the mean coefficient of variation (CV) in Mouse A (left) and Mouse B (right). The black dashed line indicates the scaling relation, which represents the theoretical expectation for statistical averaging of independent components.

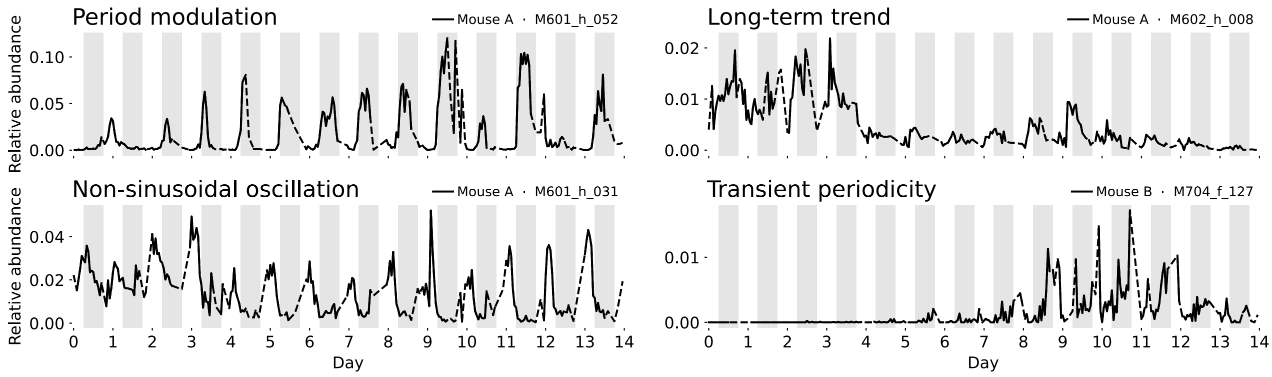


**Fig. S9** Examples of complex temporal dynamics observed in the metagenomic time-series data. Representative MAG-level time series exhibiting diverse temporal behaviors: period modulation (upper left), long-term trend (upper right), non-sinusoidal oscillation (lower left), and transient periodicity (lower right).

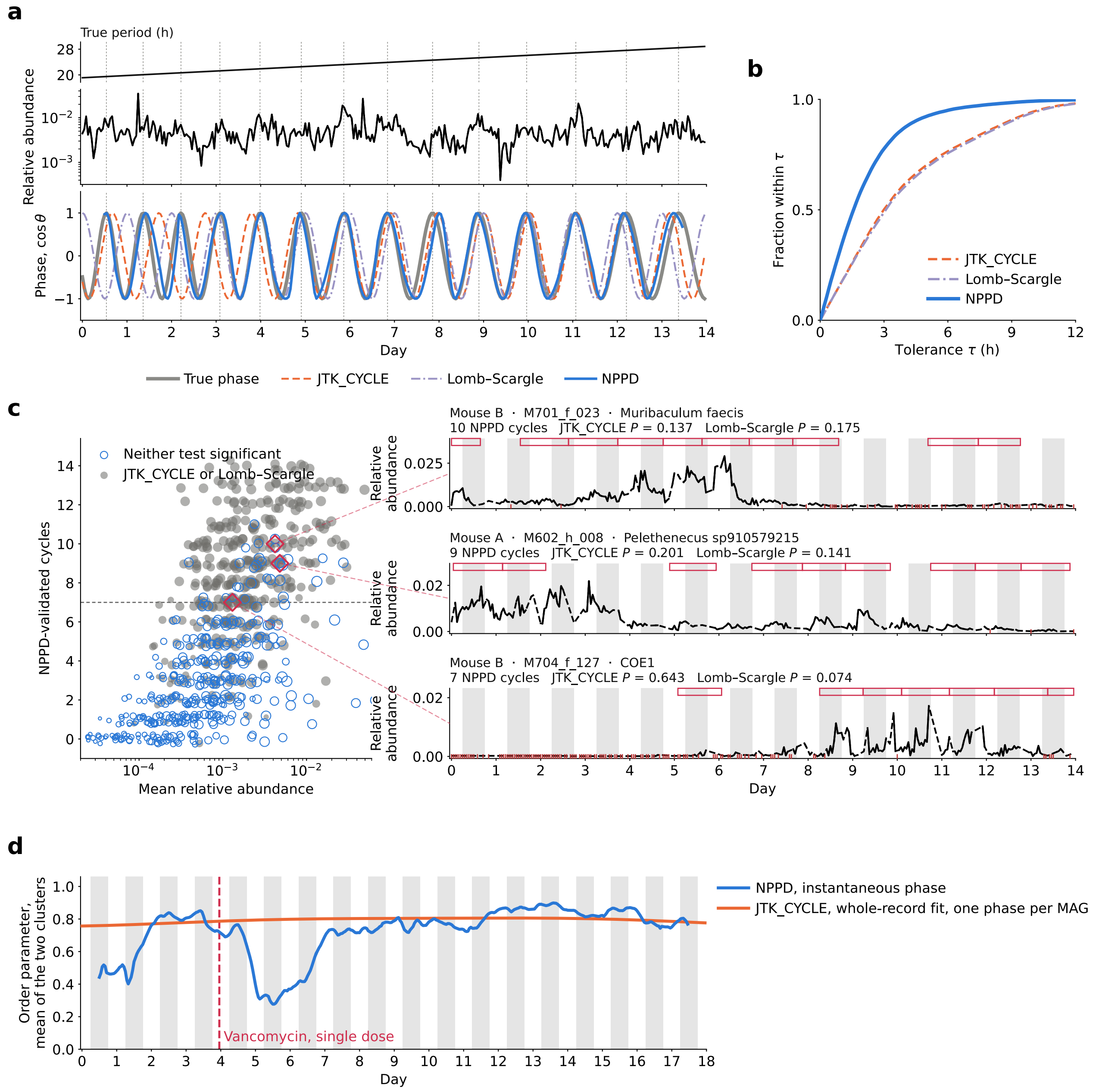


**Fig. S10** NPPD resolves time-varying local phase that is not represented by a single global period and phase. (a) Representative 14-day synthetic time series with a period that changes continuously from 19.1 to 28.8 h. Top, true local period; middle, relative abundance; bottom, true phase and phase estimates from JTK_CYCLE, Lomb–Scargle, and NPPD, shown as $\cos\theta$. A single method-specific phase offset was removed before comparison (Methods). All methods searched the same 16–32 h period range. (b) Phase-estimation accuracy across 200 simulations. Curves show the fraction of hourly estimates within tolerance $\tau$ of the true phase. Median absolute phase errors were 1.54 h for NPPD, 2.94 h for JTK_CYCLE, and 3.01 h for Lomb–Scargle. (c) Relationship between mean relative abundance and the number of NPPD-validated cycles for all 792 MAGs from Mouse A and Mouse B.

Filled grey points indicate significance by at least one global test $(q < 0.05)$; open blue points indicate significance by neither. Point size represents temporal detectability, and the dashed line indicates the seven-cycle criterion. Right, representative MAGs marked in the scatter plot; red outlines indicate NPPD-validated cycles and grey shading the dark phase. (d) Mean within-cluster phase coherence during vancomycin perturbation in Mouse D, $(R_{\text{Light}}(t) + R_{\text{Dark}}(t))/2$, calculated from the same predefined 120 MAGs. Blue, NPPD instantaneous phase; orange, phase reconstructed from a single whole-record JTK_CYCLE fit.

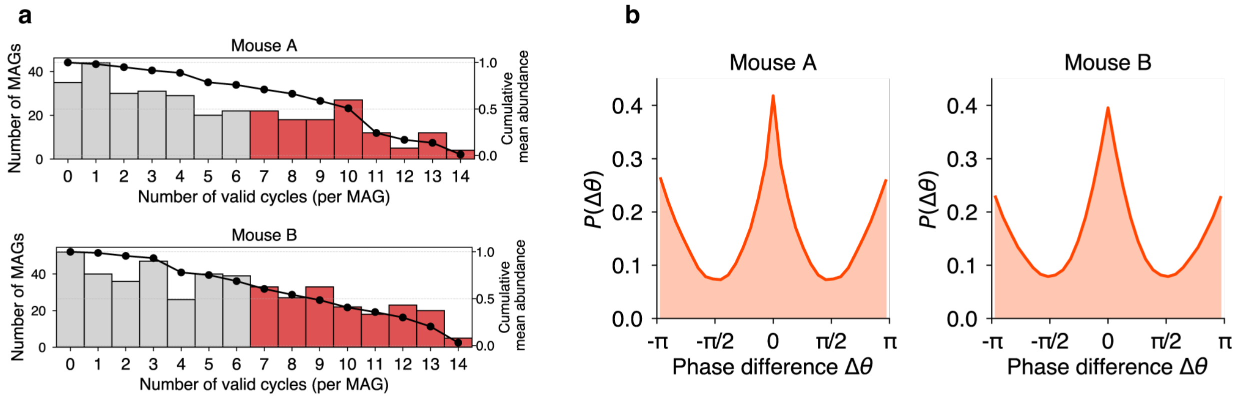


**Fig. S11** Core oscillatory MAGs and phase difference structure. (a) Histogram of valid oscillatory cycles per MAG in Mice A and B. The red bars highlight MAGs with ≥ 7 valid cycles, representing the core set used for subsequent phase-based analyses. The black line indicates the cumulative relative abundance of MAGs with at least that number of cycles. (b) Distribution of phase differences Δθ between all MAG pairs in Mice A and B, computed only from statistically significant oscillatory cycles.

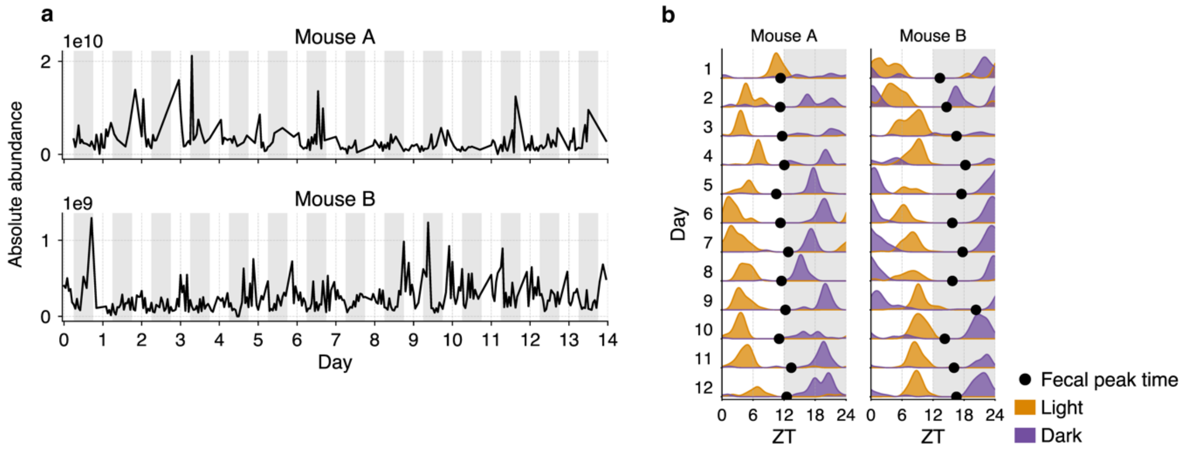


**Fig. S12** Temporal dynamics and phase behavior of absolute MAG abundances estimated using a spike-in standard. (a) Time series of total absolute MAG abundance in Mice A and B over the 14-day observation period, estimated using the *Salinibacter ruber* spike-in standard. (b) Phase behavior of the Light (orange) and Dark (purple) clusters based on absolute abundances. Daily distributions of peak times showing their temporal shifts over 14 days. Black dots indicate the timing of fecal excretion peaks.

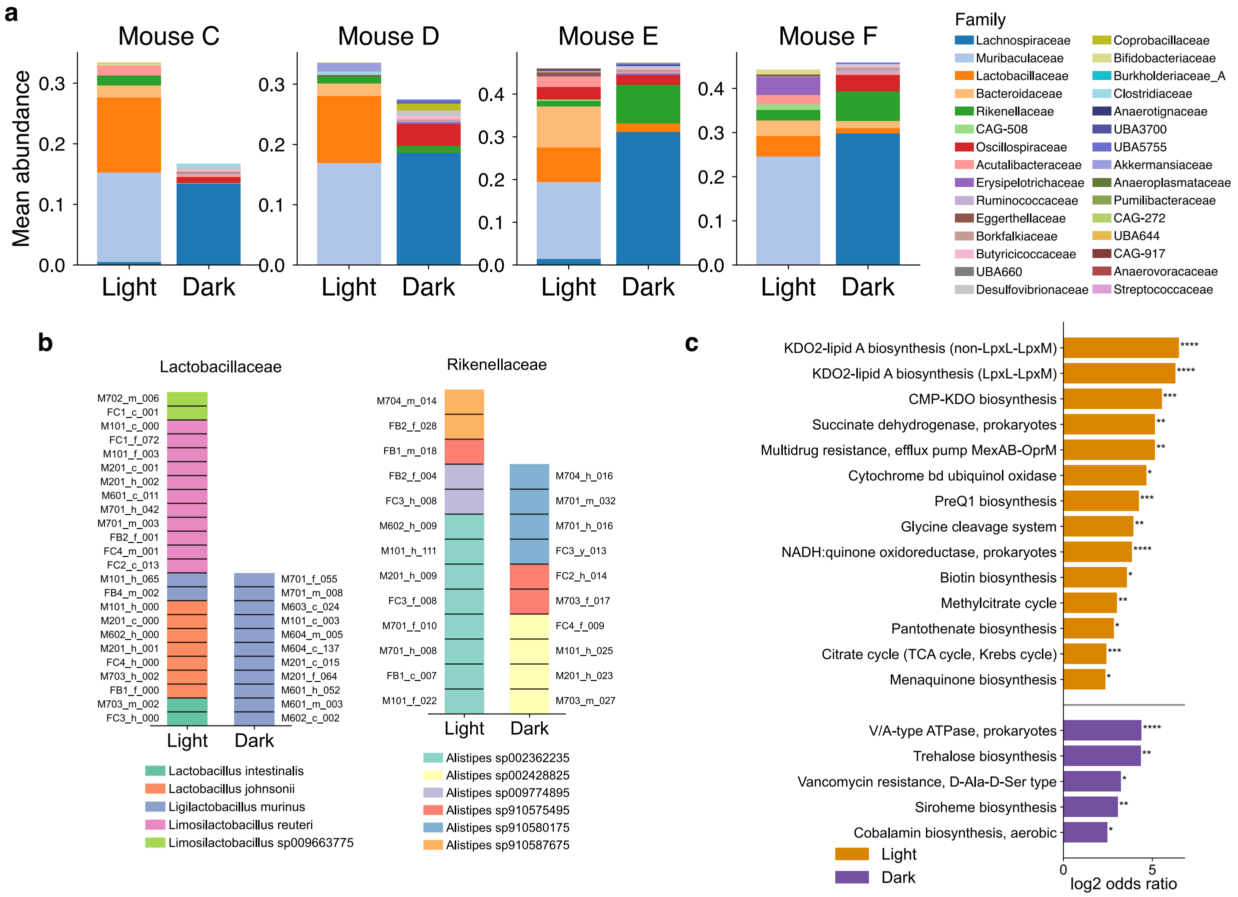


**Fig. S13** Taxonomic and functional characteristics of Light and Dark clusters across mice. (a) Mean family-level composition of the Light and Dark clusters in Mice C–F. Stacked bars indicate the relative abundance of each family within each cluster. (b) Species composition of the families Lactobacillaceae (left) and Rikenellaceae (right) in the Light and Dark clusters for Mice C–F. Stacked bars indicate the MAGs belonging to each family detected in each cluster. (c) KEGG modules significantly enriched in each cluster (Fisher's exact test, $P < 0.05$). Bars represent the $\log_2$ odds ratio for enrichment in the Light (orange) and Dark (purple) clusters.

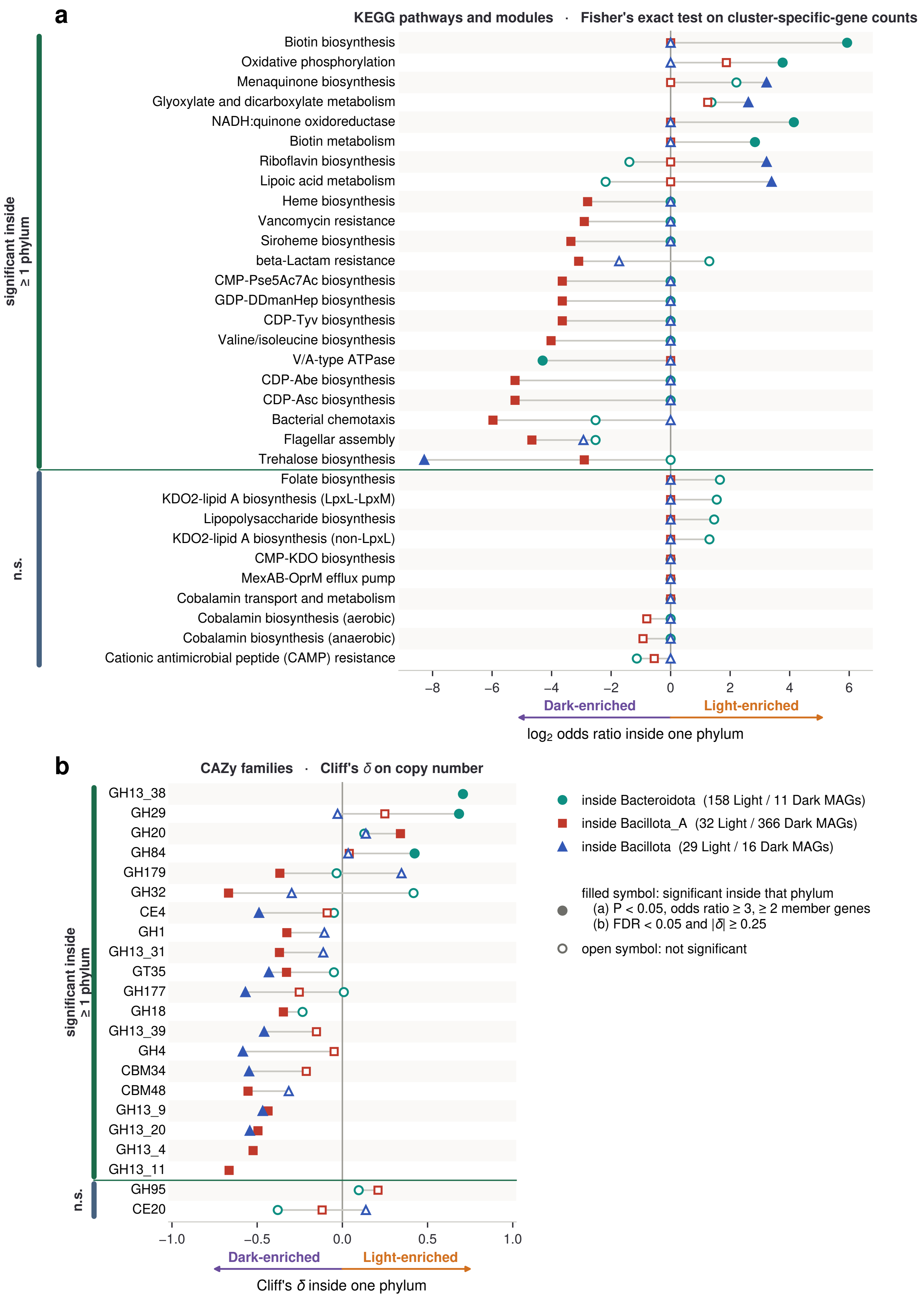

a
KEGG pathways and modules · Fisher's exact test on cluster-specific-gene counts
significant inside ≥ 1 phylum
Biotin biosynthesis
Oxidative phosphorylation
Menaquinone biosynthesis
Glyoxylate and dicarboxylate metabolism
NADH:quinone oxidoreductase
Biotin metabolism
Riboflavin biosynthesis
Lipoic acid metabolism
Heme biosynthesis
Vancomycin resistance
Siroheme biosynthesis
beta-Lactam resistance
CMP-Pse5Ac7Ac biosynthesis
GDP-DDmanHep biosynthesis
CDP-Tyv biosynthesis
Valine/isoleucine biosynthesis
V/A-type ATPase
CDP-Abe biosynthesis
CDP-Asc biosynthesis
Bacterial chemotaxis
Flagellar assembly
Trehalose biosynthesis
n.s.
Folate biosynthesis
KDO2-lipid A biosynthesis (LpxL-LpxM)
Lipopolysaccharide biosynthesis
KDO2-lipid A biosynthesis (non-LpxL)
CMP-KDO biosynthesis
MexAB-OprM efflux pump
Cobalamin transport and metabolism
Cobalamin biosynthesis (aerobic)
Cobalamin biosynthesis (anaerobic)
Cationic antimicrobial peptide (CAMP) resistance
−8 −6 −4 −2 0 2 4 6
Dark-enriched
Light-enriched
log2 odds ratio inside one phylum
b
CAZy families · Cliff's δ on copy number
significant inside ≥ 1 phylum
GH13_38
GH29
GH20
GH84
GH179
GH32
CE4
GH1
GH13_31
GT35
GH177
GH18
GH13_39
GH4
CBM34
CBM48
GH13_9
GH13_20
GH13_4
GH13_11
n.s.
GH95
CE20
−1.0 −0.5 0.0 0.5 1.0
Dark-enriched
Light-enriched
Cliff's δ inside one phylum
inside Bacteroidota (158 Light / 11 Dark MAGs)
inside Bacillota_A (32 Light / 366 Dark MAGs)
inside Bacillota (29 Light / 16 Dark MAGs)
filled symbol: significant inside that phylum
(a) P < 0.05, odds ratio ≥ 3, ≥ 2 member genes
(b) FDR < 0.05 and |δ| ≥ 0.25
open symbol: not significant

**Fig. S14** Functional comparison between the Light and Dark clusters within individual phyla. The analyses used in the main text were repeated with the comparison restricted to MAGs within each phylum, pooling the six individually profiled mice. Symbols show results for *Bacteroidota* (circles; 158 Light / 11 Dark MAGs), *Bacillota_A* (squares; 32 / 366), and *Bacillota* (triangles; 29 / 16), with grey lines connecting the same feature across phyla. Filled symbols indicate significance within that phylum and open symbols indicate non-significance. Positive values indicate enrichment in the Light cluster and negative values enrichment in the Dark cluster. (a) KEGG pathways and modules. Within each phylum, cluster-associated genes were identified using the same Mann–Whitney U test, Cliff's δ, and Benjamini–Hochberg correction as in the main analysis, and enrichment of each pathway or module was assessed by Fisher's exact test. The horizontal axis shows the $\log_2$ Haldane–Anscombe odds ratio. (b) CAZyme families. Copy numbers were compared between Light and Dark MAGs using the same two-sided Mann–Whitney U test and Cliff's δ as in the main analysis, with Benjamini–Hochberg correction within each phylum.

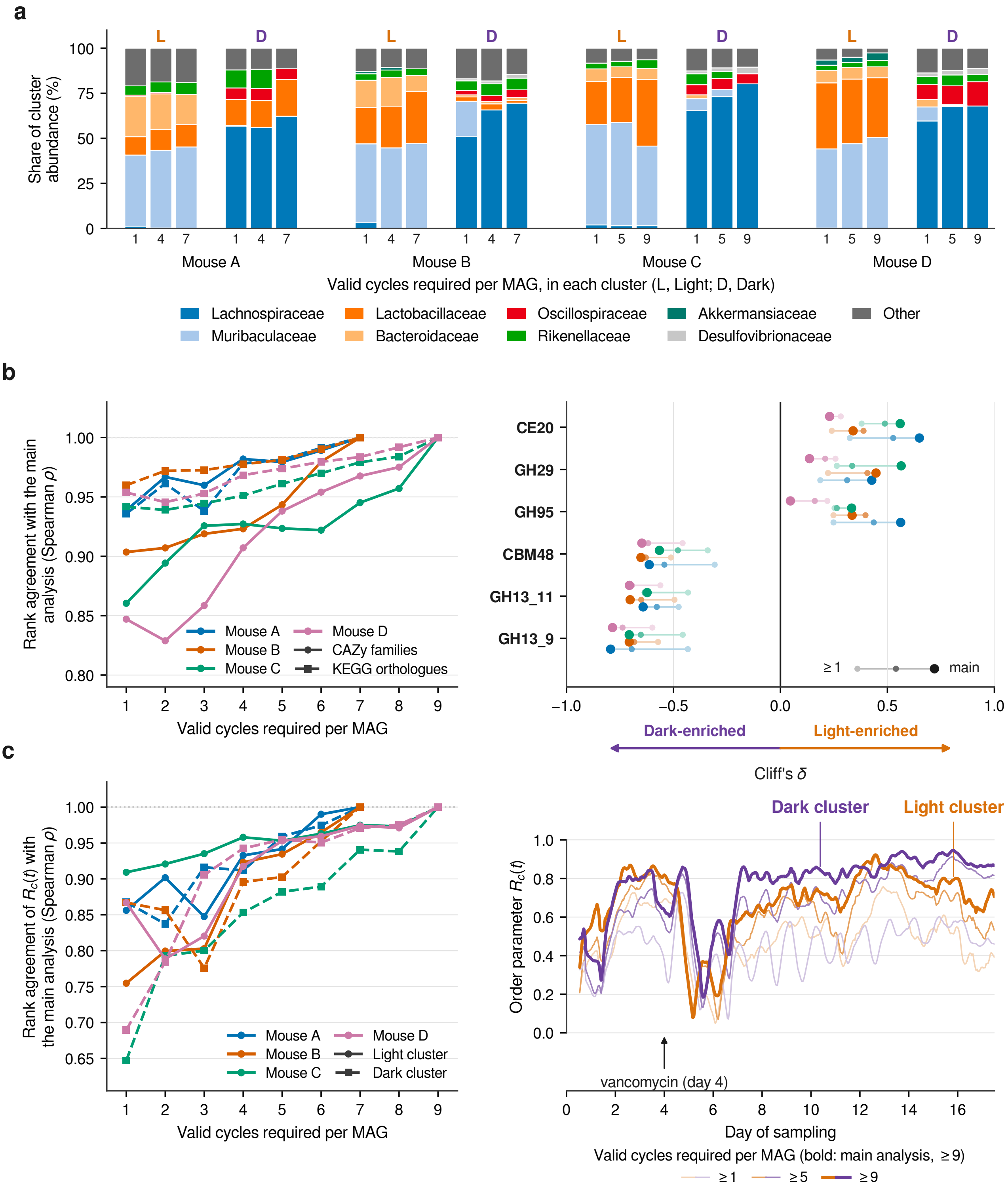


**Fig. S15** Phase-based results are robust to the valid-cycle criterion used to define core oscillatory MAGs. The criterion was relaxed stepwise from that used in the main analysis to ≥1 validated cycle, and downstream analyses were recomputed at each threshold. For panels showing selected thresholds, 1, 4, and 7 cycles are shown for the 14-day records and 1, 5, and 9 cycles for the 18-day records.

(a) Family-level composition of the Light (L) and Dark (D) clusters across thresholds (cf. Fig. 4c). The eight most abundant families are shown individually and the remainder as Other. (b) Functional contrast between the two clusters. Left, Spearman correlation of the Light-versus-Dark effect-size vector (Cliff's δ) at each threshold with that of the main analysis, for CAZy families (solid) and KEGG orthologues (dashed). Right, Cliff's δ for the CAZy families highlighted in Fig. 4f at three representative thresholds; larger points indicate the main-analysis criterion. (c) Cluster phase coherence. Left, Spearman correlation between the ¥(R_c(t)¥) trajectory at each threshold and that of the main analysis for the Light (solid) and Dark (dashed) clusters. Right, $R_c(t)$ for Mouse D at three representative thresholds; the main-analysis criterion is shown in bold and the arrow indicates vancomycin administration.

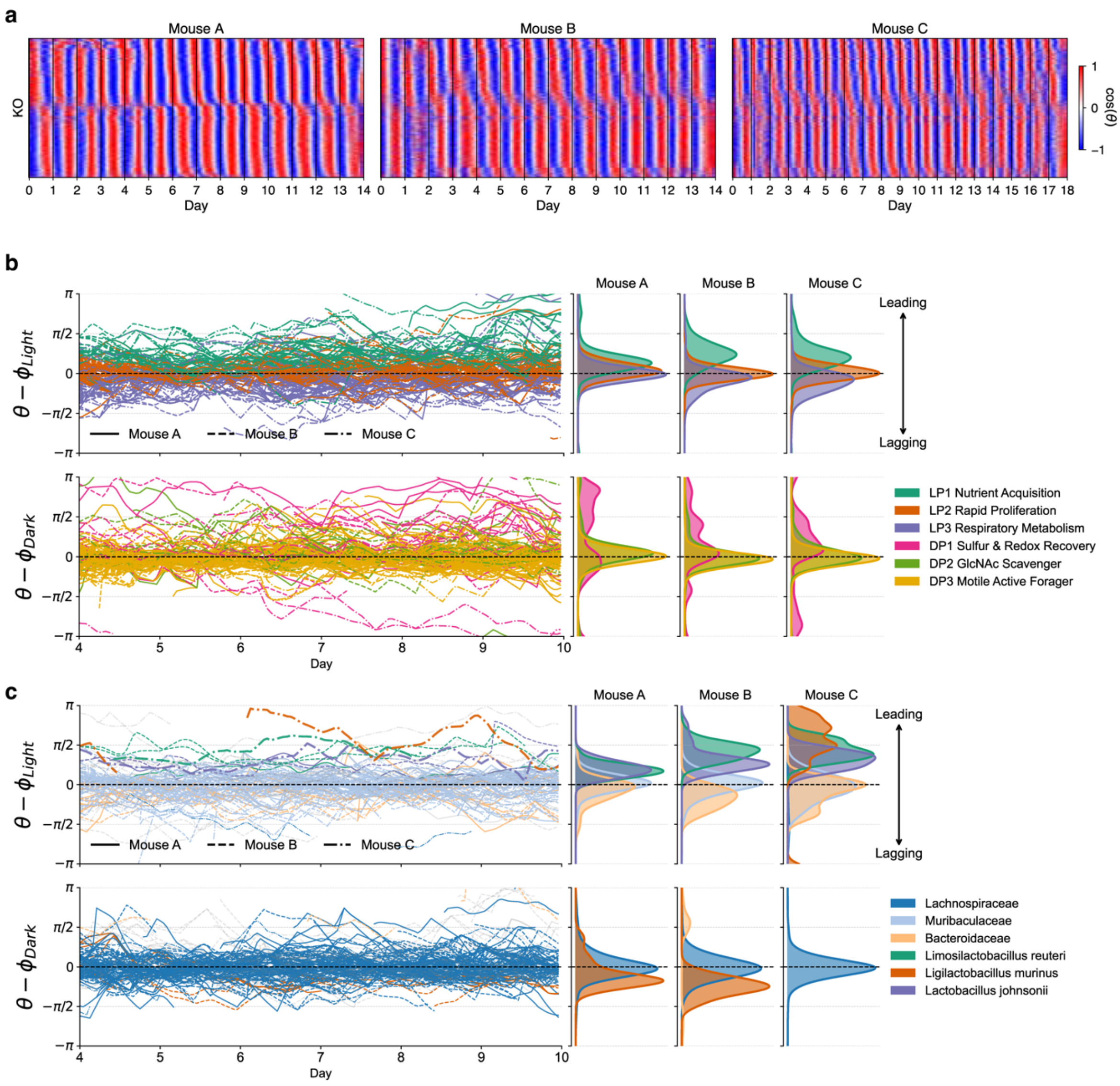


**Fig. S16** Phase organization of functional and taxonomic dynamics across mice. (a) Phase heatmap of KOs for Mice A–C. The horizontal axis represents time (days), and each row corresponds to a KO. KOs are ordered according to their mean peak time in Mouse A. (b) Relative phase dynamics of KOs grouped by functional gene subsets. Relative phase is defined with respect to the mean phase of each cluster. Time series of KOs in the Light and Dark clusters are shown together with their distributions for each mouse. Colors indicate functional gene subsets. (c) Relative phase dynamics of MAGs grouped by taxa. Time series of MAGs in the Light and Dark clusters are shown together with their distributions for each mouse. Colors indicate taxonomic groups.
